# Attentional focus and emotion modulate voice recognition deficits in cerebellar stroke patients

**DOI:** 10.64898/2025.12.08.692937

**Authors:** Leonardo Ceravolo, Emma Stiennon, Ioana Constantin, Emilie Chassot, Marine Thomasson, Jordan Pierce, Lukas Sveikata, Alexandre Cionca, Roberta Ronchi, Emmanuel Carrera, Didier Grandjean, Frédéric Assal, Julie Péron

**Affiliations:** Neuroscience of Emotion and Affective Dynamics Laboratory, Department of Psychology and Swiss Centre for Affective Sciences, University of Geneva, Geneva, Switzerland; Clinical and Experimental Neuropsychology Laboratory, Department of Psychology, University of Geneva, Geneva, Switzerland; Swiss Centre for Affective Sciences, University of Geneva, Campus Biotech, Geneva, Switzerland; Cognitive Neurology Unit, Department of Neurology, University Hospitals of Geneva, Geneva, Switzerland; Cognitive and Affective Neuroscience Lab, Department of Psychology, University of Nebraska-Lincoln, Lincoln, Nebraska 68588, USA; Medical Image Processing Laboratory, Neuro-X Institute, École Polytechnique Fédérale De Lausanne (EPFL), Geneva, Switzerland; Faculty of Medicine, University of Geneva, Geneva Medical Center, Geneva, Switzerland; Stroke Unit, Department of Neurology, University Hospitals of Geneva, Geneva, Switzerland

**Keywords:** Vocal emotion, Voice prosody, Cerebellar stroke, Attention, fMRI, Drift diffusion model

## Abstract

The cerebellum, long regarded as a motor structure, is increasingly recognized for its role in higher-order cognitive and socio-emotional functions. Its contribution to vocal emotion decoding, however, remains insufficiently understood. While prior work has linked the cerebellum to attentional control and predictive coding, direct evidence for its role in modulating prosody recognition under explicit versus implicit attentional demands is lacking.

This study investigated how cerebellar stroke—controlled for time since stroke in months, and with an equal number of left- and right-lateralized lesions—alters vocal emotion processing depending on attentional focus. We aimed to disentangle sensory encoding and integration from decisional contributions of the cerebellum by combining behavioral analyses, drift diffusion modeling (DDM), and functional MRI in cerebellar stroke.

Fifteen patients with chronic, isolated cerebellar stroke and fifteen matched controls performed two tasks on vocal stimuli expressing anger, happiness, or neutrality. In the explicit task, participants categorized the expressed emotion; in the implicit task, they categorized speaker gender while ignoring emotional tone. Behavioral performance was analyzed using mixed-effects logistic regression and DDM (angle model). Functional MRI analyses included conventional contrasts as well as model-based regressors derived from behavioral and computational parameters.

Patients showed lower accuracy than controls, with deficits particularly pronounced in the explicit task. Performance also varied by emotion: gender recognition for angry voices (implicit processing) was relatively preserved, whereas happy and neutral voices were more vulnerable. Contrary to predictions, DDM parameters did not differ between groups. Instead, task effects dominated: implicit processing yielded higher drift rates, larger boundary separation, faster non-decision times, and stronger attentional focus compared to explicit recognition, suggesting that explicit evaluation imposes additional cognitive costs. At the neural level, patients recruited extended networks during explicit processing, including orbitofrontal cortex, amygdala, anterior insula, and cerebellar lobule IX, alongside cerebello-cortical tracts. Implicit processing was associated with more restricted activations, particularly in frontal opercular and cerebellar regions (lobule IX). Model-based analyses further revealed that successful categorization in patients relied on the left inferior frontal gyrus, inferior parietal lobule, and pre-supplementary motor area, although these effects were not significant when controlling for time since stroke.

Our findings demonstrate that the cerebellum does not primarily shape decision dynamics but optimizes sensory representations of prosodic cues for downstream evaluation. When predictive tuning is compromised, patients maintain intact decision processes yet rely on compensatory cortical recruitment, particularly during explicit tasks. This pattern supports predictive coding accounts of cerebellar function in socio-emotional communication and highlights the need to consider subtle socio-affective deficits in cerebellar patients.

## Introduction

The cerebellum, traditionally associated with motor control, is increasingly recognized for its contributions to higher-level cognitive and affective functions, including attention and emotion processing^1–4^. Through its widespread connections with cortical and subcortical structures (e.g., basal ganglia, prefrontal and temporal regions), the cerebellum supports prediction and error monitoring in a variety of domains, extending to the recognition of vocal emotion processing^5,6^. Emotional prosody, defined as segmental and suprasegmental vocal cues conveying the speaker’s affective state^7^, requires the integration of low-level auditory features in superior temporal regions with higher-order evaluative processes in frontal and limbic areas^8–16^.

Over the past decade, converging neuroimaging and clinical findings have demonstrated the role of the cerebellum in vocal emotion decoding^5,16–25^. Patients with cerebellar stroke showed selective impairments in emotional attribution^26,27^, and neuroimaging has revealed cerebellar activations during prosodic and pitch-related tasks^28–30^. These results suggest that the cerebellum participates both in sensory acquisition as well as in higher-level cognitive aspects of affective voice processing, possibly via a metacognitive function^31^. However, the cognitive processes underlying the interaction between the cerebellum and both cortical and subcortical networks during emotional decoding remain poorly understood.

One critical and unresolved issue concerns the influence of attentional mechanisms, including top-down. Vocal emotion recognition can occur under explicit conditions, when attention is directed toward the affective content of the voice, or under implicit conditions, when emotion is processed automatically while attention is focused on another aspect of the voice (e.g., speaker gender)^19,32–34^. Neuroimaging studies have shown that distinct neural networks are recruited in these conditions: explicit decoding engages frontal and temporal brain regions (e.g., the inferior frontal gyrus, IFG; the anterior cingulate cortex, ACC; the superior temporal gyrus, STG), whereas implicit decoding additionally involves structures such as the basal ganglia^19,33,35^. Yet, the cerebellum has often been overlooked in these paradigms, either due to limited field of view in fMRI or to a primary focus on cortical regions. Consequently, there is no direct evidence for cerebellar modulation by attentional focus during emotional prosody processing.

Indirect findings nonetheless point to an influence of attention on the cerebellum during the processing of emotional prosody. The cerebellum is indeed strongly implicated in attentional control, including in shifting and the allocation of attentional resources^36–38^, and deficits in these processes (sometimes termed “attentional dysmetria”) have been reported after cerebellar lesions^37^. Moreover, in other modalities such as facial emotion processing, distinct cerebellar subregions appear to support implicit versus explicit decoding^39,40^. In the auditory domain, behavioral results suggest that attentional focus influences error monitoring during vocal processing^19^, consistent with the predictive coding function of the cerebellum^31,41,42^. Implicit tasks, such as identifying speaker gender when the stimulus carries emotional prosody, engage automatic, rapid, and often subcortical mechanisms involving the vermis, alongside the amygdala and anterior cingulate cortex^43^. In contrast, explicit emotional categorization tasks require conscious evaluation and recruit posterior cerebellar regions (Crus I/II) alongside frontal and temporoparietal cortical areas implicated in executive control and theory of mind^43^. These dissociable pathways suggest that cerebellar involvement in emotion perception varies depending on the attentional and cognitive demands of the task. Taken together, these data favor the viewpoint of the cerebellum as a credible hub for integrating prediction, error monitoring, and top-down modulation during affective voice recognition and/or its role in integrating emotion with sensorimotor and cognitive signals to regulate predictive coding under uncertainty.

The present study was therefore designed with the above-mentioned aspects in mind, to specifically investigate how attentional focus would modulate cerebellar activity during emotional prosody decoding in patients with chronic cerebellar stroke (>5 months post-stroke) compared to healthy, matched controls. In a paradigm testing both explicit (emotion recognition) and implicit (gender categorization) conditions, we tested whether cerebellar contributions to vocal emotion processing are differentially engaged depending on top-down attentional demands—namely, in explicit and implicit emotion processing tasks. To capture underlying decision dynamics, behavioral data were analyzed using drift diffusion modeling, with predictions of reduced decision quality (drift rate), increased response caution (boundary separation) and slower perceptual-motor processing (non-decision time)^19,39,43–47^, and decreased attentional focus (theta)^5–7,34,43,47–49^ in patients, while initial response bias (starting point) is expected to remain intact and comparable between groups. Globally, computational modeling is rarely used to study stroke, which is a gap this study fills with the hope of promoting its use in this field. Neuroimaging predictions include altered activations in frontal, temporal, and subcortical regions (e.g., inferior frontal gyrus^14,19,33,45,46,50–52^, basal ganglia^31,35,43,48,53^), with the expected additional involvement of the amygdala and orbitofrontal cortex during explicit tasks^45,47^. Key contrasts will focus on Emotion × Task × Group interactions and group differences in neural correlates of decision accuracy. By combining wholebrain fMRI, computational modeling, and a lesion approach in 15 patients and 15 matched controls, this study provides a direct test of cerebellar involvement in top-down modulation and predictive coding during vocal emotion recognition.

## Materials and Methods

### Participants

Patients with cerebellar stroke (N=15; N=8 and N=7 left- and right-lateralized lesions, respectively; see Supplementary Fig.1 for an overview of the lesions and their overlap) and control neurologically unimpaired participants (N=15) were included in the study. Controls were matched for age and education level (years of education) with the patients, and no between-group difference was observed for these variables (age: F(1,28)=1.04, *p>*.1; education: F(1,28)=0.15, *p>*.1). For the controls, mean age was 62.06 years (SD=10.10), mean education was 17.00 years (SD=3.29) and for the patients mean age was 58.19 years (SD=11.38) and mean education was 16.50 years (SD=4.08). Using G*Power3 software^54^, and considering a group-comparison approach with worse performance expected for patients compared to controls (one-tailed), we obtain a power of 56.6% with an effect size of 0.5, an alpha of 0.05 and two samples of 15 participants (N_total_=30). For the patients, the time since the cerebellar stroke at data acquisition was 26.81 months on average (SD=24.87, range: 5-89). Concerning neuropsychological tests, all participants performed the MoCA (Montreal Cognitive Assessment test) and the PEGA (Protocole d’Evaluation des Gnosies Auditives). Mean and standard deviation values were the following: a) MoCA, Controls 28.13 (SD=1.06), Patients 27.73 (SD=2.60); PEGA, Controls 28.57 (SD=1.34), Patients 28.73 (SD=1.79). For both tests, no difference was found between groups (MoCA, F(1,28)=0.30, *p>*.1; PEGA, F(1,28)=0.07, *p>*.1). Additional exclusion criteria were the presence of: a patent foramen ovale, metallic implant(s), pacemaker, intracranial implant(s)/clip(s), a copper coil for women, auditory deficits, neurodegenerative disease(s), psychiatric or other prior neurological history—including obsessive-compulsive disorder, mild cognitive impairment, multiple sclerosis, other stroke(s) outside the cerebellum. All participants/patients gave their written informed consent to take part in the study, which was accepted by the Geneva state-wide ethics committee of the Geneva University Hospital (Comité Cantonal d’Ethique en Recherche, ‘CCER’, validated application nr. 2024-00174) and conducted according to the declaration of Helsinki and local, state-wide regulations.

### Procedure

The vocal stimuli consisted of two speech-like but semantically meaningless sentences extracted from Banse and Scherer’s validated database^55^. These pseudo-sentences were spoken in a neutral(‘interest’ category in the database), happy (‘elation’) or angry (‘hot anger’) tone by two female and two male actors, resulting in a total of 24 different stimuli. Among them, 18 stimuli were used in the present study and assigned to each task/block (Fig.1A-C), keeping the balance between actor gender and emotions. In the present study, these binaurally recorded auditory stimuli were played through MRI-compatible headphones and displayed using E-Prime (http://www.pstnet.com/eprime.cfm) in two consecutive runs (Fig.1**A**). Participants therefore performed a categorization task with two levels of the Task factor (implicit, explicit) and three for the Emotion factor (anger, happiness, neutral). See Fig.1**B** for more details. This paradigm was divided into 16 counterbalanced blocks (6 trials per block) per run, leading to a grand total of 192 trials per participant. During the eight ‘implicit’ blocks, participants had to discriminate the gender of the stimulus. For the other eight blocks, the explicit processing was evaluated by the capacity of discriminating the emotional content of the stimuli. The participants had to indicate whether the stimulus was ‘happiness’ or ‘neutral’ / ‘anger’ or ‘neutral’ by pressing the corresponding key. They did not know prior to the stimulus whether they would be presented with happy and neutral or angry and neutral choices. For all blocks, the order of the emotion presented was pseudo-randomized to avoid repetitions of the same emotion several times in a row and to avoid boundary effects (alternating emotion as the last trial of each block).

**Figure 1:**
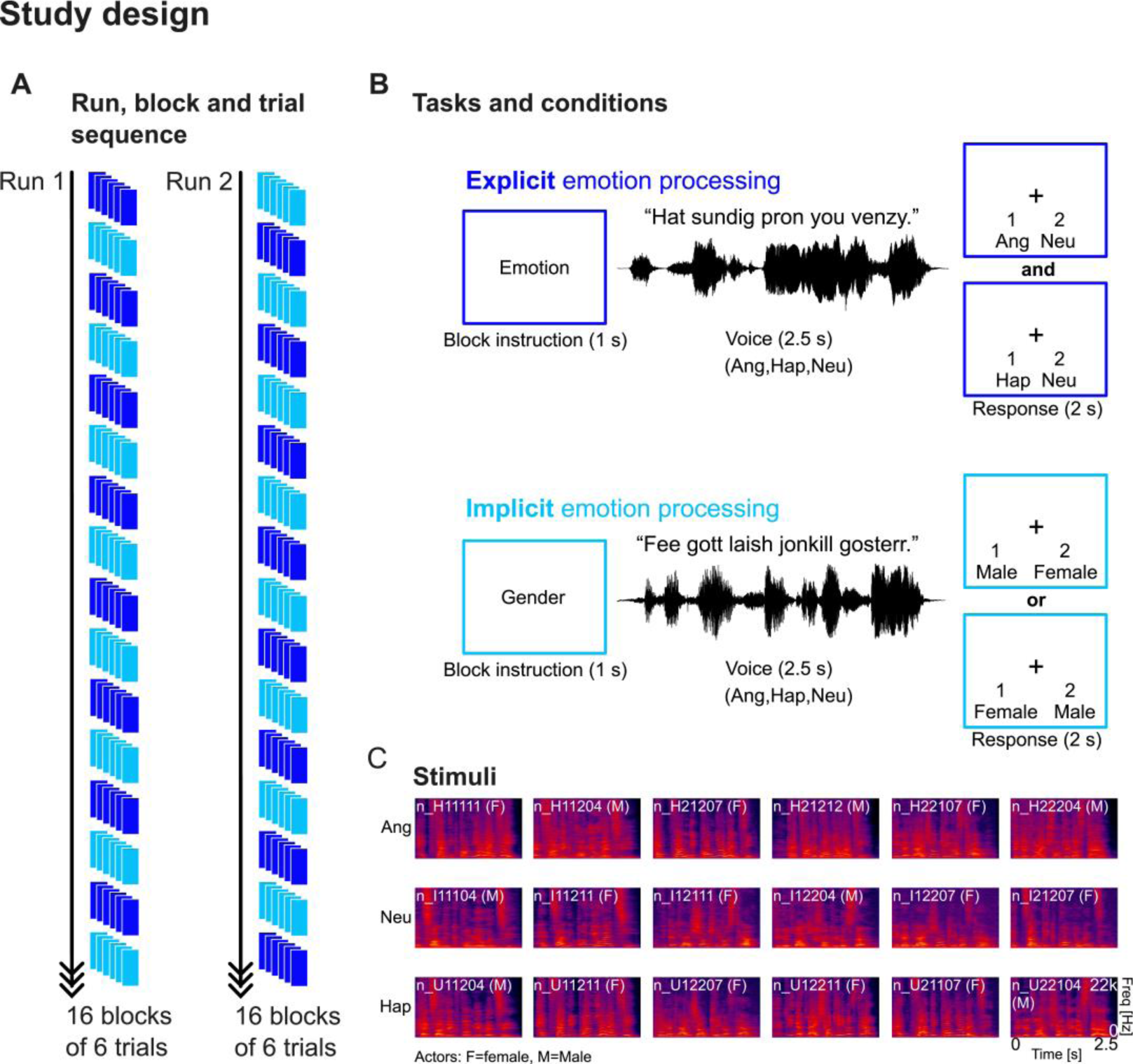
Study design, task and stimuli. (**A**) Tasks were split into two runs of equal length (16 blocks of 6 trials each), alternating between the explicit and the implicit vocal emotion processing tasks (dark and light blue, respectively). Tasks are described in (**B**), starting with an instruction of the type of block to come, then the stimulus was presented for 2.5 s and the participants were instructed to categorize these according to the task (2 s). (**C**) Illustration through a spectrogram of the 18 stimuli used in the study, with stimulus identity and speaker gender (F: female, M: male). Ang: anger; Hap: happiness; Neu: neutral.

We assumed that during the gender decision task, participants would focus on the gender of the voice, while the emotional tone of the voice would still be processed implicitly. Before each block, a message indicating “Gender” or “Emotion” was displayed on the screen for 1 s as an instruction for the upcoming block. The vocal stimuli were then played (duration: 2.5 s, see Fig.1**C**) while a fixation cross was presented at the center of the screen, then the two possible responses were displayed on the screen for 2 s. Each of the two runs lasted approximately 10 minutes.

### Behavioral data analysis: regression analysis

Behavioral results illustrate the probability of accurately recognizing voice gender or emotion –namely, the Task factor (implicit, explicit anger block, explicit happy block), as a function of the Emotion (angry, happy or neutral voice prosody) and Group (controls, cerebellar stroke patients) factors. These analyses were computed using mixed effects, regression analyses for responses (of-interest) and reaction times (of-no-interest) using R studio^56^. Complete data aggregation yielded to a matrix of 5614 lines for all participants, including Patients and Controls. The data matrix was then cleaned by excluding trials with reaction times faster than 200 milliseconds and also by excluding the 5% most extreme values of the data based also on reaction times (95^th^ percentile). This cleaning process reduced the matrix to 5201 lines/data points (7.35% of data excluded in total), with an average trial count per participant of 173.

For the ‘response’ dependent variable, we used a logistic regression using the lme4 ‘glmer’ package^57^ with voice Emotion interacting with Task and Group as fixed effects. Random effects included the random slopes of the participants (ParticipantID) with Emotion and the time since stroke (TSS; in months)—namely ‘1+TSS+Emotion|ParticipantID’, the random intercept of participant gender (ParticipantGender), stimuli (Stimulus), runs (Run), stimulus order (StimulusOrder), voice actor identity (ActorID) and actor Gender (ActorGender). The model formula was the following:

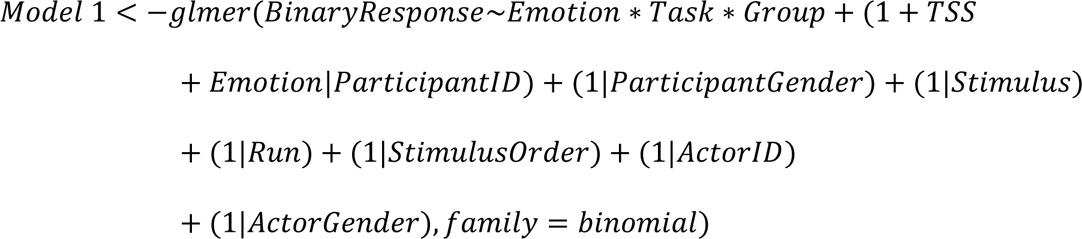

For the ‘reaction times’ dependent variable, we used a linear regression using the lme4 ‘lmer’ package with the exact same variables and formula as above, namely:

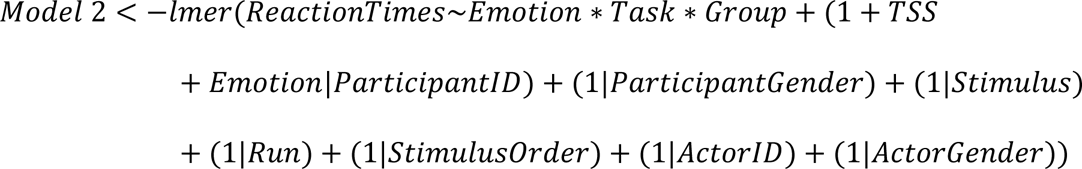

For our planned comparisons, we used a standard confidence interval of 95% and a Bonferroni correction for multiple comparisons. Specific contrasts were computed using the ‘emmeans’ (Estimated Marginal Means, or Least-Squares Means) package^58^. Further details are reported in the supplementary data.

### Computational analysis of response and reaction times data using drift diffusion modeling

We turned to computational modeling of the data to consider them in the framework of decision models, namely drift diffusion^59–61^ in the context of our study (Drift Diffusion Modeling, ‘DDM’). We chose this modeling because it is adequate for two-alternative forced-choice tasks. It models decision-making as a process that accumulates evidence until a threshold is reached, at which point a choice is made. Among the parameters of DDM, we were particularly interested in drift rate (speed of evidence accumulation, ‘*v*’, arbitrary units) and boundary separation (the threshold that separates the two choices, ‘*a*’, arbitrary units), as these parameters are critical to explain both reaction times and accuracy and could highlight fundamental decision-making processes between our patients and control participants. In more specific terms, drift rate illustrates the average speed, and direction of evidence accumulation. Therefore, a higher drift rate indicates fast evidence accumulation and potentially faster and more accurate decisions. Boundary separation illustrates the amount of evidence required to reach a certain decision: wider boundary separation translates a greater need for evidence, leading to more cautious, slower, and potentially more accurate decision-making; conversely, smaller boundary separation suggests faster, but more error-prone or impulsive, decisions. Additional and default computed parameters included ‘starting point’ (initial level of evidence before accumulation, ‘*z*’, arbitrary units), ‘non-decision time’ (accounting for processes that are unrelated to evidence accumulation, for instance the motor response, ‘*t*’ or also ‘*ndt*’, in seconds) and theta (‘θ’) for the angle model (representing the angle, in degrees, with which decision boundaries collapse over time). All aspects relating to the toolbox used (Hierarchical Sequential Sampling Modeling or ‘HSSM’ 0.2.5, running in Python) and all specific parameters and settings, analysis pipeline and cross-validation are reported in the supplementary data and in Supplementary Fig.2.

### MRI data

#### MRI data acquisition

All functional imaging data were recorded on a 3-T Siemens Prisma-fit System (Siemens, Erlangen, Germany) scanner equipped with a 32-channel antenna. A 3D sequence was used to acquire a high-resolution T1-weighted image (1mm isotropic voxels; TR=1900ms; TE=2.27ms; matrix resolution=256 x 256; FA=9degrees; 192 total slices).

For the task, functional images were acquired in descending order using a multi-band echo-planar imaging sequence of 54 slices aligned along the anterior-posterior commissure (voxel size: 2.5mm isotropic; slice thickness=2mm; TR=1300ms; TE=20ms; FOV=205 x 205mm; matrix resolution=84 x 84; FA=64degrees; BW=1952Hz/px). Functional image acquisition was continuous throughout each run of the task.

#### MRI data analysis

For each group, data were preprocessed using SPM12 (SPM12, Wellcome Trust Centre for Neuroimaging, London, UK), the CONN toolbox^62^ and the Artifact detection tools (ART) toolbox (https://www.nitrc.org/projects/artifact_detect/). Functional data were first converted to 4D Nifti in SPM12 for each run separately to get one single file per run, for more efficient computation performance and to decrease storage volume. Preprocessing steps included the following: realignment and unwarping, slice-timing correction, artifact detection followed by an additional denoising (scrubbing including functional regression and functional bandpass [0.01-0.1Hz]). At this stage, direct segmentation and normalization into the Montreal Neurological Institute (MNI) space^63^ was performed to allow comparison between participants and with other existing fMRI studies. Finally, data were spatially smoothed with an isotropic Gaussian filter of 8 mm full width at half maximum. Following the creation of a participant-specific matrices including trial onset and reaction times for each run, preprocessed, smoothed functional images were then analyzed with SPM12 both at the participant-level (first-level) as well as at the group-level (second-level). Seven general linear models (GLMs) were used to compute first-level statistics, all convolved with the Hemodynamic Response Function (HRF). Across all models, six head motion parameters were included as regressors of no-interest to account for movement-related variance, and time since stroke (in months) was included as a second-level covariate in all group analyses (no interaction with factor(s) and no centering, since controls all have a value of ‘0’). Trials in which an evaluation error was made for each task were also included as a concatenated regressor of no-interest. In the first modeling, regressors of interest were used to compute simple contrasts for each participant, leading to separate main effects of Angry, Neutral and Happy voices for each task and group, for a total of six simple contrasts for controls and for patients (Implicit task: Angry, Neutral and Happy voices; Explicit task: Angry, Neutral and Happy voices). This model yielded two flexible factorial second-level analyses, with factors Task, Emotion and Group. We also had the Participants factor (Factor 1 in each analysis) to consider interindividual variability, with data independence set to ‘yes,’ variance to ‘unequal.’ The same settings were used for the Group factor, while for the within factors of interest data independence was set to ‘no’, variance to ‘unequal’. Second-level models focused on between-groups differences for: the Task factor (Second-level analysis 1) and the interaction between Task and Emotion (Second-level analysis 2, Task * Emotion). The aim of this analysis was to highlight brain regions that correlate with correct voice categorization in general, between our groups. This fitted probability was obtained through the analysis of behavioral data by taking the trial-wise output of the mixed-effects generalized linear modeling of the response data and fitting it, again trial-wise. Behavioral data analysis is described in detail above. This third second-level analysis therefore focused on between-groups correlates of general voice categorization accuracy, A model-based approach was also employed in 5 additional models, using the mean posterior estimate of each of the angle DDM computational model parameters described above. Therefore, model 3 included drift rate (*v*) as parametric modulator for each trial, model 4 included boundary separation (*a*), model 5 included starting point (*z*), model 6 included non-decision time (*t*) and model 7 included the theta parameter (θ). These five models were computed to respectively investigate neural correlates of decision efficiency and quality, response caution, initial response preference or bias, perceptual-motor processing speed and finally attentional focus or selectivity. For all of these five model-based analyses, we then used a two-sample t-test second-level analysis (Second-level analyses 3-7), with only the contrast images of the parametric modulator, across Run, Task and Emotion but per Group. Independence between samples (Controls, Patients) was set to ‘yes’ and variance was set to ‘unequal’.

Activations of all seven second-level models were ultimately thresholded in SPM12 (https://www.fil.ion.ucl.ac.uk/spm/software/spm12/) by using a voxel-wise FDR correction at *p*<.05 and an arbitrary cluster extent of k>10 voxels per cluster—to remove extremely small clusters and/or single voxel activations. Thresholded contrast activations were then rendered on brain templates from the CONN toolbox^62^. Regions were labelled using the latest version of the ‘automated anatomical labelling’ (‘aal3’) atlas^64,65^.

## Results

### Behavioral measures and computational modeling

Behavioral data included as dependent variables were response accuracy (Fig.2) and reaction times (Supplementary Fig.3; of no-interest). Detailed results values for these dependent variables are summarized in Supplementary Table 1.

**Figure 2.**
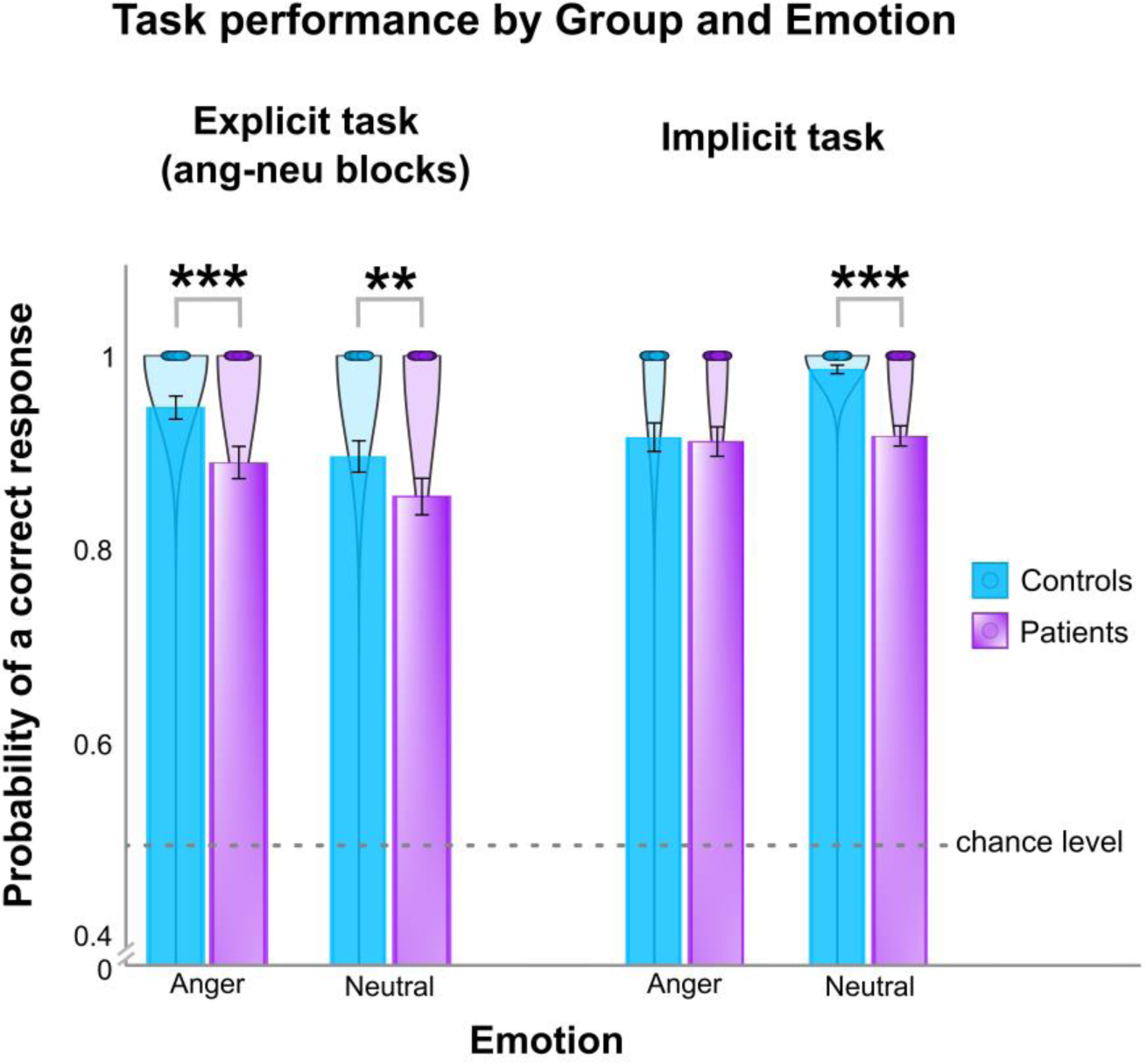
Probability of correct voice categorization as a function of Emotion and Task—including block types, for both groups. Probability of accurate voice categorization (*y* axis) for the explicit and implicit tasks in both control participants (blue) and cerebellar stroke patients (purple) and according to each emotion except happiness (Anger/Neutral, *x* axis). Bars represent the mean values, error bars the standard deviation from the mean. Chance level of 50%, as indicated by the horizontal, dashed grey line. ** *p*<.01; *** *p*<.001.

#### Probability of a correct response (response accuracy)

Response data of each task and for each group were treated as ‘correct’ or ‘incorrect’ (coded as ‘1’ or ‘0’). Logistic regression was used to interpret these data as the probability of a correct response as a function of our Task, Emotion and Group factors (fixed effects) in addition to relevant random effects (intercepts and slopes, including time since stroke in months as a function of Group, see Methods).

As mentioned in the methods, we report the outcomes of both between- and across-block types for the explicit task. The modeling reported in this section therefore includes splitting the explicit task as a function of the existing architecture of the two types of blocks (anger blocks, happiness blocks). For this model, R^2^m=37.83% and R^2^c=58.87%. See Fig.2 for the detailed graphical representation of the results. Since both controls and patients were at or below chance level to classify happy and neutral voices in the happy-neutral block types of the explicit task, no comparisons were computed with this block type and we also do not interpret happy voices in the implicit task, for the sake of coherence (the modeled values are still reported in Supplementary Table 1). We therefore only tested our hypothesis for angry and neutral voices for both tasks, although all emotions were modeled in the analysis.

By splitting blocks of the explicit task by their type, we observed no general effect of voice Emotion (χ^2^(2)=0.69, *p*>.10), but a main effect Task (including block types, χ^2^(2)=411.76, *p*<.001) and Group (χ^2^(2)=23.68, *p*<.001). Two-way interactions revealed interacting Emotion and Task type factors (χ^2^(2)=10.38, *p*<.01) and Task and Group (χ^2^(2)=9.72, *p*<.01) but only a tendency towards significance for the interaction between Emotion and Group (χ^2^(2)=4.82, .05<*p*<.1). The three-way interaction of interest between Emotion, Task and Group was also significant (χ^2^(2)=13.91, *p*<.001).

To test our specific hypothesis concerning an impact of cerebellar stroke on voice categorization both in explicit and implicit situations and for angry, happy and neutral voices, we computed planned comparisons based on the significant three-way interaction. Namely, we contrasted our groups for each task and emotion—except for happy voices, as explained above. Comparing groups for each task and emotion resulted in the following differences: cerebellar stroke patients performed worse than matched controls for angry (β=1.22, 95% CI [0.55, 1.90], z=3.54, *p*<.001) and for neutral (β=0.82, 95% CI [0.22, 1.42], z=2.69, *p*<.01) voices in the explicit task and for neutral voices in the implicit task (β=2.21, 95% CI [1.46, 2.96], z=5.76, *p*<.001), while they interestingly performed similarly to controls for angry voices in the implicit task (β=0.36, 95% CI [-0.29, 1.00], z=1.08, *p*>.10). See Fig.2.

#### Computational data modeling using ‘angle’ DDM, integrating responses and reaction times

Throughout this section and in the discussion, we will refer here to each parameter as a function of its primary role in the context of angle DDM as well as in psychological machinery. Therefore, drift rate (*v*) illustrates decision efficiency and quality, boundary separation (*a*) translates response caution or speed-accuracy tradeoff, starting point (*z*) illustrates an initial response preference or bias, non-decision time (*t*) represents perceptual-motor processing speed and finally theta (θ) represents attentional focus or selectivity. As mentioned in the methods, these parameters are estimated by integrating accuracy and reaction time data on a trial-by-trial basis. See Supplementary Table 2 for the detailed main effects comparisons and Bayesian statistical values, with posterior probability (‘pp’) instead of p-values, and Supplementary Table 3 for the two-way and three-way interactions (the data revealed no evidence of any interaction effects).

The computational data revealed inconclusive effects, for all parameters and for the Population factor, meaning that controls and patients did not globally differ in decision efficiency, caution, initial response preference, executive functions or attentional focus (mean difference for: *v*=-0.026, pp=.465; *a*=0.014, pp=.505; *z*=0.017, pp=.535; *t*=0.015, pp=.525; θ=0.024, pp=.600, respectively). The results of the parameter estimates go against our hypotheses according to which patients would develop compensatory mechanisms due to cerebellar stroke. Finally, strong evidence was observed for the Task factor, for all parameters but starting point (*z* mean difference=-0.009, pp=.30). Therefore, independently of Group or Emotion, the implicit compared to the explicit task revealed better decision efficiency and quality (*v* mean difference=1.191, pp=1.00), increased evidence accumulation (*a* mean difference=0.234, pp=1.00), faster voice processing and response execution (*t* mean difference=-0.036, pp=.00) and finally a stronger attentional focus on task-relevant features (θ mean difference=0.117, pp=1.00). All other comparisons can be found in Supplementary Table 2. This is in line with the mixed-effects behavioral data analyzed with a logistic regression and reported above.

### Neuroimaging results

Following behavioral data analyses and computational modeling, we then tested our effects of interest in accordance with our hypotheses—namely the three-way Emotion * Task * Group and two-way Task * Group interactions. We therefore describe here: a) wholebrain neuroimaging data analysis of traditional group-level contrasts; b) model-based modeling of the data illustrating the general neural underpinnings of the probability of correct voice categorization between groups; c) ‘angle’ DDM computational modeling with parameter-associated results. For the analyses presented in this section, the time since stroke (in months) was used as a second-level control covariate (no interaction with factor(s), no centering).

#### Wholebrain data for Task and Group

The interaction between Task and Group was of the highest interest in the present study. We were especially interested in the patients > controls comparison, in order to investigate the impact of cerebellar stroke on the explicit and implicit categorization of vocal emotion and its neural underpinnings. For the explicit task, comparing patients to controls revealed a widespread cortical and cerebellar network of brain regions (Fig.3; Supplementary Table 4). In the cortex, we observed enhanced activity in the primary somatosensory and motor cortices as well as in the pre-supplementary motor area, superior temporal gyrus and sulcus and anterior insula (Fig.3**A-C**). We also observed a large cluster of activity in the ventromedial prefrontal cortex, more specifically in the medial (Fig.3**D,F**), mid and superior (Fig.3**E**) orbitofrontal cortex, amygdala and parahippocampal gyrus (Fig.3**D,F**). Subcortical activity was increased in the left caudate nucleus (Fig.3**G**), as well as in the cerebellum, especially in fiber tracts linking this region with the cortex (inferior cerebellar peduncle and internal capsula; Fig.3**H**) and in bilateral lobule IX of the cerebellum (Fig.3**I**).

**Figure 3:**
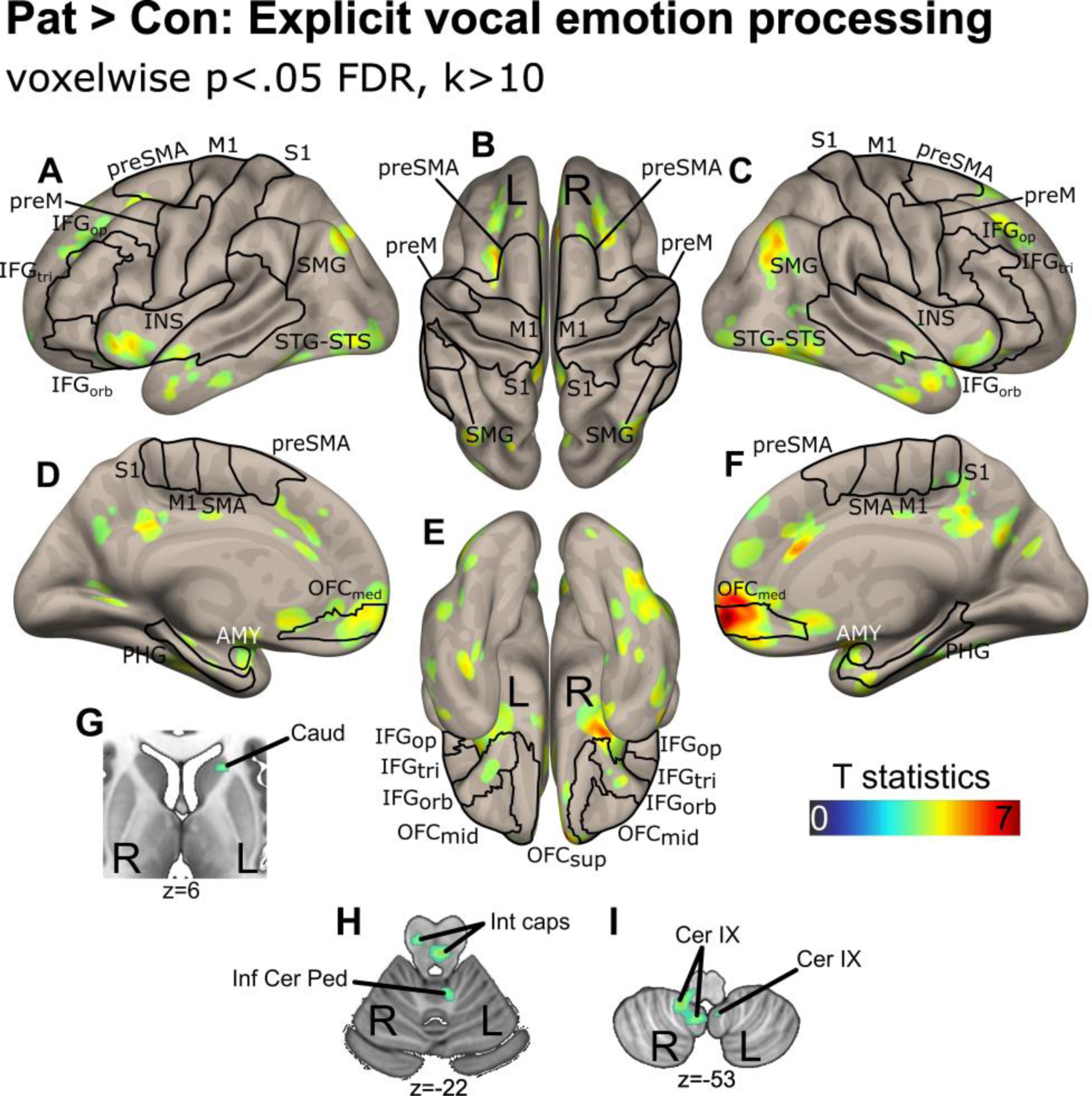
Neural activations when categorizing vocal emotion for the Explicit task, for patients compared to controls. Sagittal (**A,C**), superior (**B**), medial (**D,F**), inferior (**E**) views in addition to axial slices of the basal ganglia (**G**) and cerebellum (**H-I**) for the explicit categorization of emotional voices for patients (Pat) compared to control (Con) participants. Data corrected for multiple comparisons using voxelwise p<.05 FDR, with minimum cluster size of 10. The colorbar represents the voxelwise T value. IFG: inferior frontal gyrus; S1: primary somatosensory cortex; M1: primary motor cortex; preM: premotor cortex; SMG: supramarginal gyrus; STG/STS: superior temporal gyrus/sulcus; INS: insula; OFC: orbitofrontal cortex; AMY: amygdala; SMA: supplementary motor area; PHG: parahippocampal gyrus; Caud: caudate nucleus; Thal: thalamus; Inf Cer Ped: inferior cerebellar peduncle; Int Caps: internal capsule; Cer: cerebellar lobule; Ver: vermis. Suffixes: op: *pars opercularis*; tri: *pars triangularis*; orb: *pars orbitalis*; med: medial; sup: superior. L/R: left/right hemisphere.

Examining activation that was greater for matched controls than patients in the explicit task was less informative regarding our hypotheses but for the sake of exhaustivity we computed this contrast, that revealed STG/STS and inferior frontal gyrus activations in addition to cerebellar activation limited to a small cluster in left Crus I area. See Supplementary Fig.4. The triple interaction between Emotion * Task * Group was not significant, and no voxels survived voxel-wise FDR thresholding.

When contrasting patients to matched controls for the implicit task, we observed left inferior frontal activations—in the *pars opercularis*, left-lateralized premotor and pre-supplementary motor area (preSMA) activations in addition to left anterior insula and precuneus activations (Fig.4**A-C,D,F**). In subcortical regions, no enhanced activity was observed in the basal ganglia or amygdala (Fig.4**D-F**), but we observed increased activity in the cerebellum, more specifically in left lobule IX (Fig.4**G**). See Supplementary Table 5 for detailed clusters coordinates.

**Figure 4:**
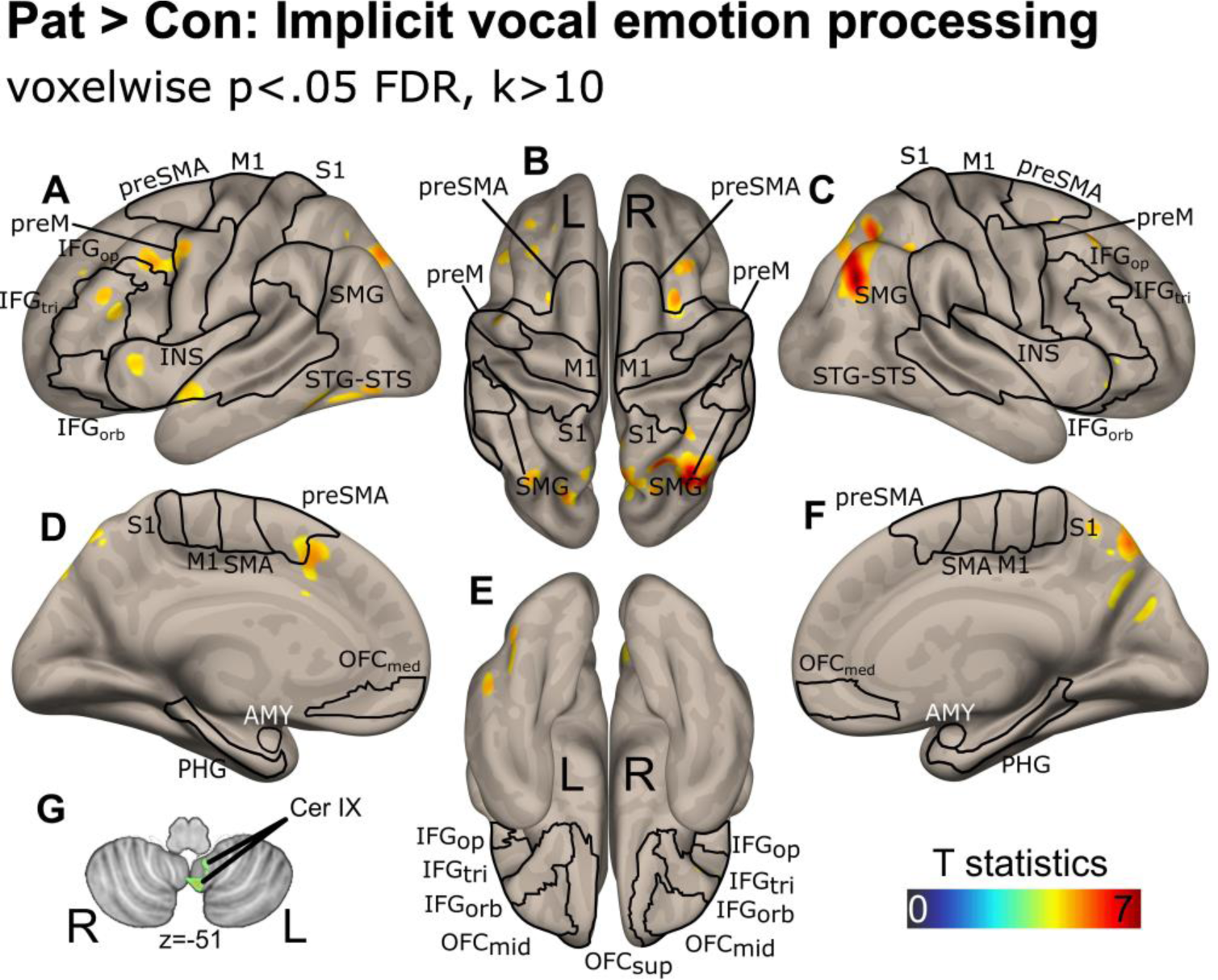
Neural activations when categorizing vocal emotion for the Implicit task, for patients compared to controls. Sagittal (**A,C**), superior (**B**), medial (**D,F**), inferior (**E**) views in addition to an axial slice of the cerebellum (**G**) for the implicit categorization of emotional voices for patients (Pat) compared to control (Con) participants. Data corrected for multiple comparisons using voxelwise p<.05 FDR, with minimum cluster size of 10. The colorbar represents the voxelwise T value. IFG: inferior frontal gyrus; S1: primary somatosensory cortex; M1: primary motor cortex; preM: premotor cortex; SMG: supramarginal gyrus; STG/STS: superior temporal gyrus/sulcus; INS: insula; OFC: orbitofrontal cortex; AMY: amygdala; SMA: supplementary motor area; PHG: parahippocampal gyrus; Cer: cerebellar lobule. Suffixes: op: *pars opercularis*; tri: *pars triangularis*; orb: *pars orbitalis*; med: medial; sup: superior. L/R: left/right hemisphere.

Comparing controls to patients for this task did not yield to any voxels surviving the corrected voxelwise statistical threshold. Similarly, no activations survived corrected statistical thresholding for the triple interaction between Emotion, Task and Group.

#### Brain correlates associated with the probability of correct voice categorization across tasks, by Group

To take a step further and examine whether brain regions are directly associated with the probability of correctly categorizing the emotional voice stimuli across tasks and between groups, we employed model-based fMRI data analysis. The general mechanism underlying the probability of a correct explicit (per block type) and implicit categorization in patients compared to controls did not reveal any above-threshold voxels when controlling for time since stroke. However, not controlling for this second-level covariate revealed very specific left-lateralized brain enhancement in the cortex but not in any subcortical structures. More specifically, we observed the recruitment of the left inferior frontal gyrus—*pars triangularis* mainly but also *pars opercularis*, inferior parietal lobule and the pre-supplementary motor area. These brain areas therefore represent the functional neural underpinnings of the general mechanism relied upon when cerebellar stroke patients need to explicitly and implicitly categorize vocal emotions, compared to matched control participants, but only if we don’t consider the time since the cerebellar stroke of the patients. See Supplementary Fig.5.

#### Brain correlates associated with the parameter estimates of DDM angle modeling of the data

As mentioned before, drift diffusion modeling of the data using the ‘angle’ distribution was computed. Except for initial response bias (*z*), we expected differences between our groups for all other parameters (drift rate ‘*v*’, boundary separation ‘*a*’, non-decision time ‘*t*’ and theta ‘θ’) but we did not find any such effects both behaviorally and for neuroimaging data. We did, however, find results common to both Controls and Patients for drift rate (Fig.5, Supplementary Table 6), boundary separation (Supplementary Fig.6) and theta (Supplementary Fig.7). Controls and cerebellar stroke Patients share common brain regions underlying decision quality and efficiency (drift rate), decision caution or speed-accuracy tradeoff (boundary separation) and attentional focus (theta).

**Figure 5:**
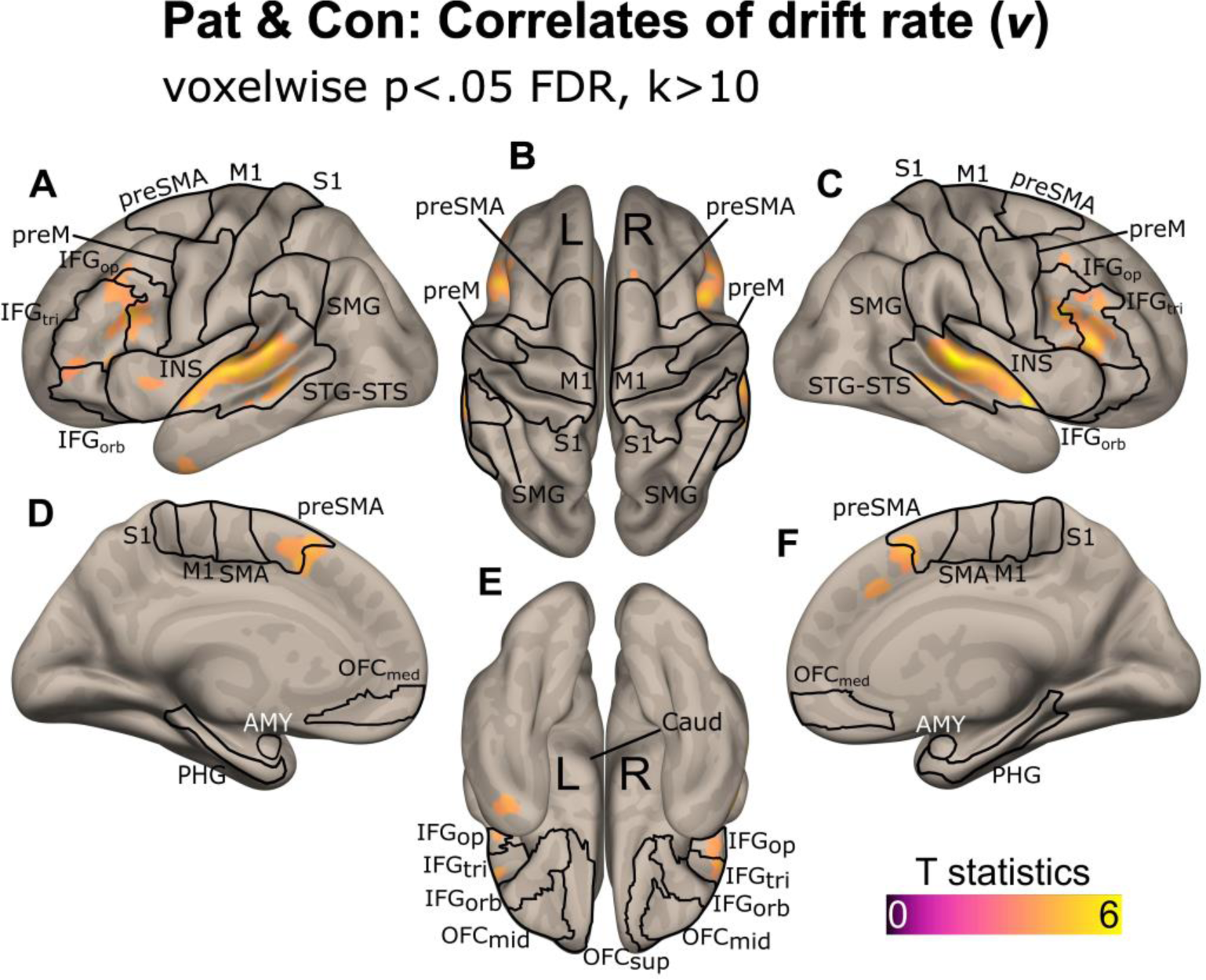
Neural correlates of drift rate (*v*) following angle DDM modeling of the data, across Group. Sagittal (**A,C**), superior (**B**), medial (**D,F**), inferior (**E**) views for decision quality and efficiency, as represented by angle DDM drift rate for Patients (Pat) and Controls (Con). Data corrected for multiple comparisons using voxelwise p<.05 FDR, with minimum cluster size of 10. The colorbar represents the voxelwise T value. IFG: inferior frontal gyrus; S1: primary somatosensory cortex; M1: primary motor cortex; preM: premotor cortex; SMG: supramarginal gyrus; STG/STS: superior temporal gyrus/sulcus; INS: insula; OFC: orbitofrontal cortex; AMY: amygdala; SMA: supplementary motor area; PHG: parahippocampal gyrus. Suffixes: op: *pars opercularis*; tri: *pars triangularis*; orb: *pars orbitalis*; med: medial; sup: superior. L/R: left/right hemisphere.

Drift rate was especially relevant, with decision quality and efficiency correlating with bilateral superior temporal brain areas such as the STG, but also with subparts of the IFG and left anterior insula (Fig.5**A,C**). Further correlates included the most medial and anterior portion of the preSMA (Fig.5**D,F**), while no subcortical regions correlated with drift rate.

Overlaps were found in the bilateral superior temporal and inferior frontal gyrus as well as in the preSMA with correlates of decision caution (boundary separation parameter) and to a lesser extent with attentional focus represented by the theta parameter (see Supplementary Fig.6 and Supplementary Fig.7, respectively).

## Discussion

In this study, we provide evidence that cerebellar lesions selectively impair vocal emotion recognition, with effects modulated by both attentional demands and emotion category—when accounting for the time elapsed since stroke. These findings once again extend the role of the cerebellum beyond motor and attentional control, highlighting its contribution to the optimization of sensory representations underlying affective evaluation. Our results suggest that the cerebellum fine-tunes the precision of prosodic input signals, thereby facilitating their integration within fronto-temporo-limbic networks. When this tuning mechanism is disrupted, as in cerebellar stroke, accuracy decreases despite preserved decision dynamics, and compensatory recruitment of broader cortical networks becomes necessary. Taken together, these findings favor a closer examination of the behavioral signatures associated with cerebellar damage, their computational underpinnings, and the neural mechanisms that allow patients to maintain some level of performance despite impaired sensory optimization.

### Behavioral effects

Consistent with existing literature^66–68^, patients with cerebellar lesions showed overall lower accuracy compared to controls across tasks. Crucially, group differences were modulated by task demands and task structure, with larger deficits emerging during explicit emotion recognition, in line with the proposed role of the cerebellum in top-down attentional control^2–4,19,32^. Interestingly, the effects also varied by emotion category. In the implicit task, patients were impaired for neutral but not for angry voices. In contrast, patients showed significant deficits for angry and neutral voices in the explicit task. Because happy voices were recognized below-chance level by both groups in the explicit emotional task, we refrained from making any interpretation involving happy voices across both tasks. We think, however, that the ‘interest’ category of the voices in the original validation study—that we used as a ‘neutral’ category here—was probably the source of this unexpected observation, especially when happiness was relatively low in the stimuli, thus confusing participants. Overall, the above-mentioned pattern of results indicate that the cerebellum may differentially contribute to the processing of discrete emotional categories, possibly depending on their salience or motivational relevance. The preservation of anger processing in the implicit task may reflect the automatic prioritization of threat-related signals by limbic circuits. However, when attentional resources are explicitly required (i.e., in the explicit task), cerebellar damage seems to hinder this processing, leading to the observed impaired recognition.

Contrary to our predictions, drift diffusion modeling did not reveal systematic differences between patients and controls in decision parameters. This result suggests that core decision computations (evidence accumulation, response caution, and attentional selectivity) remain relatively intact after cerebellar stroke for higher-level judgement. Instead, strong and consistent task effects were observed for both groups taken together: the implicit task was associated with more efficient decision-making (higher drift rate), increased evidence accumulation (higher boundary separation), faster processing (shorter non-decision time), and stronger attentional focus (higher θ) compared to the explicit task. This effect is coherent with the mixed-effects modeling of the behavioral responses mentioned above, for which we observed a higher probability of accurate emotional voice recognition in the implicit than in the explicit task, especially for angry voices. It is also in line with existing literature on automatic attentional capture by emotional prosody^5,69,70^ or studies in which prosody is processed explicitly^50^. Since a control for several additional random factors was not feasible with angle DDM to maintain a satisfactory level of model convergence—see methods and supplementary methods, we can therefore assume that random effects and especially slopes and/or task design may have played a role in the absence of angle DDM interactions with the Group factor. These findings indicate that explicit attention to emotional content imposes additional cognitive demands, making decision processes less efficient irrespective of lesion status. At the emotional level, angry voices were consistently associated with reduced decision efficiency, in line with prior work showing that negative emotions, and particularly threat-related signals, impose higher perceptual and decisional costs^19,33,35^.

The discrepancy between the behavioral group differences and the absence of computational parameter differences suggests that cerebellar stroke may primarily impair sensory representations (e.g., encoding and integration of prosodic cues) rather than the decisional processes per se—this interpretation is also backed by the absence of between-groups cognitive difference for the MoCA. In this view, the cerebellum would contribute to optimizing the input signal for cortical and subcortical evaluative processes, consistent with predictive coding accounts of cerebellar functions^31,41,42^. Lesions in the cerebellum would therefore reduce the ‘quality’ of the input, lowering accuracy, but without necessarily altering the internal dynamics of evidence accumulation once the decision process has been initiated. In other words, the observed pattern of computational results suggests that commonalities still exist in the neurocognitive evaluation—namely, in the decisional aspects—of voice emotions both in cerebellar stroke patients and in matched control participants at the behavioral level.

### Neural correlates of emotional voice processing after cerebellar stroke

Wholebrain functional neuroimaging results complement the behavioral findings by showing that cerebellar stroke patients rely on an extended cortical and subcortical network when categorizing emotional voices. During the explicit task, patients exhibited widespread enhanced activations encompassing sensorimotor, temporal, and prefrontal areas, as well as limbic structures including the orbitofrontal cortex, amygdala, parahippocampal gyrus, and bilateral anterior insula. Subcortical regions such as the caudate nucleus were also recruited, alongside cerebellar regions (lobule IX, bilaterally) and cerebello-cortical tracts. This large-scale pattern, repeatedly reported in explicit prosody studies^6,31,43^, likely reflects compensatory engagement of executive, attentional, and affective circuits to support effortful explicit decoding of vocal emotion. This result is also in line with literature on post-stroke brain plasticity^71,72^. By contrast, the implicit task mostly elicited more focal activity in the *pars triangularis* of the inferior frontal gyrus^6,33,45,51^, the preSMA, cerebellar lobule IX and the left angular gyrus, without any engagement of limbic structures. This more circumscribed activation pattern aligns with preserved behavioral performance in implicit processing, particularly for threat-related signals, and supports the view that implicit emotion recognition relies on rapid, automatic pathways that can partially bypass compromised cerebellar computations and associated regions.

Interestingly, our findings consistently implicated cerebellar lobule IX during both implicit and explicit prosodic processing. Although this region is traditionally associated with oculomotor and vestibular functions, converging recent evidence suggests that lobule IX may also play a broader role in attentional and cognitive regulation. Resting-state connectivity studies have shown that the caudal portion of lobule IX participates in the dorsal attention network^73^, positioning it as a potential node involved in the maintenance and orienting of attention. Complementarily, microstructural work indicates that integrity of cerebellar regions encompassing lobule IX predicts faster reaction times and more efficient cognitive performance^74^, consistent with a contribution to perceptual efficiency or vigilance. Although task-based fMRI studies have primarily localized cerebellar contributions to the dorsal attention network within lobules VIIb/VIIIa^75,76^, our findings raise the possibility that lobule IX may interact with these networks by supporting predictive tuning or multisensory alignment during emotional prosody processing. This interpretation aligns with models that conceptualize the cerebellum as optimizing internal predictive representations rather than directly shaping decision dynamics.

Model-based fMRI was computed to provide a more direct yet generalizable link between behavior and neural dynamics. While the fitted successful categorization probability across tasks in patients did not reveal any significant activations when controlling for time since stroke, excluding this covariate revealed the recruitment of the left inferior frontal gyrus—*pars opercularis* and *triangularis*, inferior parietal lobule, and preSMA. These regions are known to support voice-based decision-making^6,19,45,51^, executive functions such as rule maintenance^77,78^, attentional control^79,80^, and inhibitory processes^44,46^. They may thus constitute the neural scaffolding that supports performance when cerebellar contributions to sensory encoding and integration are reduced^81^. Interestingly, these activations were lateralized to the left hemisphere. Since these data are no longer significant when including time since stroke as group-level covariate, it tells us that the heterogeneity concerning the delay between stroke onset and data acquisition should not be overlooked, as it introduces added variance sufficient to dilute the impact of these brain regions on vocal emotion processing.

Finally, computational, drift diffusion modeling results also converge towards this interpretation. Despite the clear behavioral group differences with general, mixed-effects regression models, no systematic group differences were observed in decision-related parameters. Both patients and controls shared common neural correlates for drift rate, boundary separation, and attentional selectivity, including widespread bilateral STG, IFG, anterior insula and preSMA regions. This pattern indicates that the computations underlying evidence accumulation and decisional caution remain relatively preserved after cerebellar stroke, and that the observed behavioral impairments most likely originate from sensory encoding and integration rather than decision-making *per se*.

### Functional implications

Taken together, the results of our study suggest a dissociation between sensory and decisional contributions to vocal emotion recognition following ischemic cerebellar stroke—controlled for lesion lateralization. The cerebellum appears to support the optimization of sensory representations of prosodic cues, consistent with predictive coding accounts whereby it refines the precision of incoming signals for downstream evaluative processes. However, once evidence accumulation begins, decisional mechanisms unfold normally, as indicated by intact DDM parameters and shared neural correlates in patients and controls.

This interpretation also explains the emotion-specific patterns observed behaviorally. The relative preservation of implicit anger processing likely reflects the privileged and automatic processing of threat-related signals within limbic circuits—even independently of attention, which are less dependent on cerebellar precision-tuning. By contrast, explicit tasks require sustained attentional allocation and controlled evaluation, processes that heavily rely on intact cerebello-cortical loops and are therefore disproportionately affected by cerebellar damage. These results have clear implications for the patients’ everyday life, especially for social interactions and emotion comprehension—and regulation. But our data also highlight the impact of such lesions and subsequent emotion recognition difficulties when patients need to reintegrate their work environment, especially when their job(s) requires high social and conversational implication. In this case, a higher cognitive load may be needed to process such social stimuli as well as a higher need for multitasking. Our data therefore have implications for research but also for clinical aspects, especially related to rehabilitation.

### Limitations

Several limitations should be considered when interpreting these findings. First, the relatively small sample size (N=30; N=15 per group) somewhat limits statistical power, particularly in detecting more subtle or distributed effects in neuroimaging analyses—although we carefully used very conservative statistical thresholds and correction for multiple comparisons. It is very difficult to recruit patients with isolated cerebellar stroke ischemic lesions as in our study, and stroke studies, especially in functional designs, are generally underpowered for that reason. Linked to this aspect is the heterogeneity among lesion sites and size that cannot be controlled for completely, leading to inter-individual differences among patients in our study. Second, the cross-sectional nature of the study prevents differentiation between stable compensatory mechanisms and early recovery-related adaptations following cerebellar stroke—as revealed by model-based results when considering or excluding the time since stroke as a covariate. Third, although the acoustic stimuli were standardized, subtle differences in prosodic intensity or spectral features across emotional categories may have influenced perceptual and attentional demands. Fourth, while the relative fit of the computational angle DDM model was ‘good’, caution is still warranted and a more compatible study design would be highly beneficial—especially concerning the architecture of each block for each task. Finally, causal approaches such as longitudinal designs or neuromodulatory techniques (e.g., transcranial stimulation^82^) may be particularly relevant for directly probing compensatory mechanisms after stroke and should be carefully addressed in future studies.

### Conclusion

Our results demonstrate that cerebellar stroke selectively impairs vocal emotion recognition, with deficits depending on task demands and emotion category. While core decision-making mechanisms appear preserved, the cerebellum may play a key role in optimizing sensory representations of prosodic cues, thereby facilitating efficient and accurate socio-emotional evaluation. Neuroimaging revealed that patients compensate for degraded sensory precision by recruiting extended cortical and limbic networks, particularly during explicit processing. These results support predictive coding accounts of cerebellar function and underline the importance of cerebello-cortical loops in socio-emotional communication. Beyond advancing theoretical models, this work highlights the need to consider socio-affective deficits in cerebellar patients and provides a basis for developing targeted interventions to improve emotional communication after stroke.

## Data availability

Data and codes will be made available on the free repository YARETA, which meets the FAIR guidelines, through direct and permanent DOI URL upon acceptance of the manuscript for publication.

## Acknowledgements

We warmly thank the volunteers, the patients, the Hospital employees and caregivers and the team of the Brain and Behavioral Laboratory of the University of Geneva, Switzerland, for their help with setting-up MRI sequences and with MRI data acquisition.

## Funding

The present research was supported by Swiss National Science Foundation (SNSF) grants to JP (PI) and FA (Co-PI) within the project “Influence of top-down mechanisms on cerebellar activity during vocal emotion decoding” (2023-2026), Grant N°: 105314_215015 through the University of Geneva. LS was supported by Alzheimer’s Association research grant (AACSF-22-922907). The funders had no role in data collection, discussion of content, preparation of the manuscript, or decision to publish.

## Competing interests

The authors report no conflict of interests.

## Notes

### Competing Interest Statement

The authors have declared no competing interest.

