## Supplementary methods, figures and tables for "Attentional focus and emotion modulate voice recognition deficits in cerebellar stroke patients"

### Supplementary material

#### Methods

##### *Generalized mixed-effects regression*

Gender significantly improved the modeling of the data (model with Gender vs. without Gender: $\chi^2(1)=9.32$, $p<.01$, AIC=4060.3 vs. 4059.7) and was therefore added. Since control participants were matched to the cerebellar stroke patients for age and educational level, these variables were not included in the modeling of the data to avoid redundancy. We however made sure no significant difference was observed (controls vs. patients for age: $F(1,28)=1.04$, $p>.1$; controls vs. patients for education: $F(1,28)=0.15$, $p>.1$). See the 'Participant' section of the Methods for the average values of each category and each population.

##### *Computational modeling of the data: toolbox, parameters, settings*

In order to compute this type of modeling, we used the latest available version (0.2.5) of the very recent and state-of-the-art, Python-based Hierarchical Sequential Sampling Modeling (HSSM) module (https://lnccbrown.github.io/HSSM/). The HSSM module is derived from its predecessor, Hierarchical Drift Diffusion Model or 'HDDM'[1,2]. Compared to HDDM, HSSM benefits from a modern Bayesian backend, namely PyMC[3] that yields to better model convergence, faster sampling and also more reliable posteriors.

For the modeling, behavioral data were exported per Population, Participant, Task (including block type), Run, Emotion and Trial in their original order of presentation for each participant and vertically concatenated. Stimulus order was also reported as well as the trial-level binary response and related reaction time. Complete data aggregation yielded to a matrix of 5614 lines for all participants, including Patients and Controls. The matrix was then cleaned by excluding trials with reaction times faster than 200 milliseconds and also by excluding data according to a 95 percentile. This cleaning process reduced the matrix to 5201 lines/data points (7.35% of data excluded in total), with an average trial count per participant of 173. The clean data were then passed on to HSSM modeling using several relevant model distributions (the standard 'drift diffusion' model; the derived 'angle' model; the 'Ornstein-Uhlenbeck' model or 'O-U'). Settings included the 'safe' priors to avoid issues during sampling and the likelihoods were estimated using the 'approx_differentiable' option, therefore directing HSSM to use LAN (Likelihood Approximation Network) pre-trained neural networks to approximate the likelihoods. This choice enabled fast gradient-based sampling—much faster than traditional methods—by replacing costly computations (partial differential equation) with differentiable neural network approximations. Although very fast, this procedure allows for highly reliable and stable data sampling. For all models, each parameter was modeled using the following global formula:

$$y \sim 1 + \text{Task} + \text{Emotion} + \text{Population} + (1 + \text{Emotion}|\text{Population})$$

This formula was designed to model the independent main effects of the Task, Emotion and Population factors in addition to the slope of voice emotion categorization as a function of each population (Emotion|Population). For each model, data were eventually inferred from the model: data were drawn from the 'nuts_numpyro' PyMC sampler using 5 chains, 5201 draws and 1000 burn-in samples. Models—namely, DDM, angle and O-U—were then compared using Pareto smoothed importance sampling leave-one-out cross-validation (LOO) method[4,5] and the angle model was the best one, with an expected log pointwise predictive density (ELPD) score of -1131.29 with a standard error of the ELPD (dSE) of 0. This means the 'angle' model had the highest out-of-sample predictive fit, namely, the ELPD value closest to zero. The standard 'DDM' model obtained an ELPD score of -1328.86 (dSE=21.57) and the 'O-U' model -1354.47 (dSE=23.30), therefore ranking second and third, respectively. Model convergence for the DDM angle was also very good, with R-hat values way below 1.01 for all parameters and associated effects computed using the above global formula (global R-hat range across all parameters: [1.0002, 1.0004]; see Supplementary Table 8).

We therefore used Bayesian statistics and tested all the parameters ($v$, $a$, $z$, $t$, θ, see Supplementary Table 9) of the 'angle' model with our factors, namely Population, Task, Emotion and according to our hypotheses, these data are reported in the results section. To validate our angle model, we used posterior predictive checks, confirming that simulated data from the fitted model closely matched the observed distributions of reaction times and accuracy. To do so, we created 500 samples for each trial/data point (N=5201) for a total of 2'600'500 simulated data points (5201 * 500), therefore testing our fitted DDM angle model against the entire posterior distribution. This procedure was carried out by passing our model to the Sequential Sampling Model Simulators (version 0.8.3, https://github.com/lnccbrown/ssm-simulators) through HSSM. See Figure 2 below for a detailed visual comparison of observed and simulated data for both accuracy and reaction times data. Reaction times were simulated well (Supplementary Fig.2**A-F**; no difference between observed vs simulated, p=.988, Supplementary Fig.2**K-L**) with a scaling factor of 2.05 (they were over-estimated by a constant value but the pattern and proportions were respected compared to observed data; Supplementary Fig.2**L**). On the other hand, accuracy data were trickier to simulate (Supplementary Fig.2**G-J**), especially for the Task factor (Supplementary Fig.2**H**), leading to a significant under-estimation of simulated compared to observed data (observed, mean accuracy=0.823; simulated, mean accuracy=0.559, $\text{difference}_{\text{Sim-Obs}}$=-0.264, $p$=.026; Supplementary Fig.2**L**). Noteworthy is the fact though that observed and simulated accuracy still correlated strongly, with r=.785 (Supplementary Fig.2**J**).

#### Results

*Mixed effects logistic regression: artificially balanced modeling of the data*

*Reaction times data*

Although of less interest in the present study, reaction times data of each task were analyzed using a linear regression similar to the one used for response patterns, as a function of the Task, Emotion and Population factors (fixed effects, see Methods). Variance explained by the fixed effects was 3.41% ($R^2c$) while the full model including random effects ($R^2c$) explained 28.81% of the variance of the reaction times data. See Supplementary Fig.4 and Table Supplementary 1.

Regarding main effects, reaction times were mainly predicted by the Task ($\chi^2(1)$=18.05, $p$<.001) and the Population ($\chi^2(1)$=11.46, $p$<.001) factors, with generally slower reaction times for cerebellar patients than matched control participants. Only one significant—two-way—interaction was observed for Emotion * Task ($\chi^2(2)$=11.69, $p$<.01). No other effects were observed (all $p$>.1), and we therefore did not pursue any post-hoc contrasts since: i) significant effects did not include the Population factor; ii) the triple interaction was not significant; iii) reaction times were of no real interest in the present study. See Supplementary Fig.3 below.

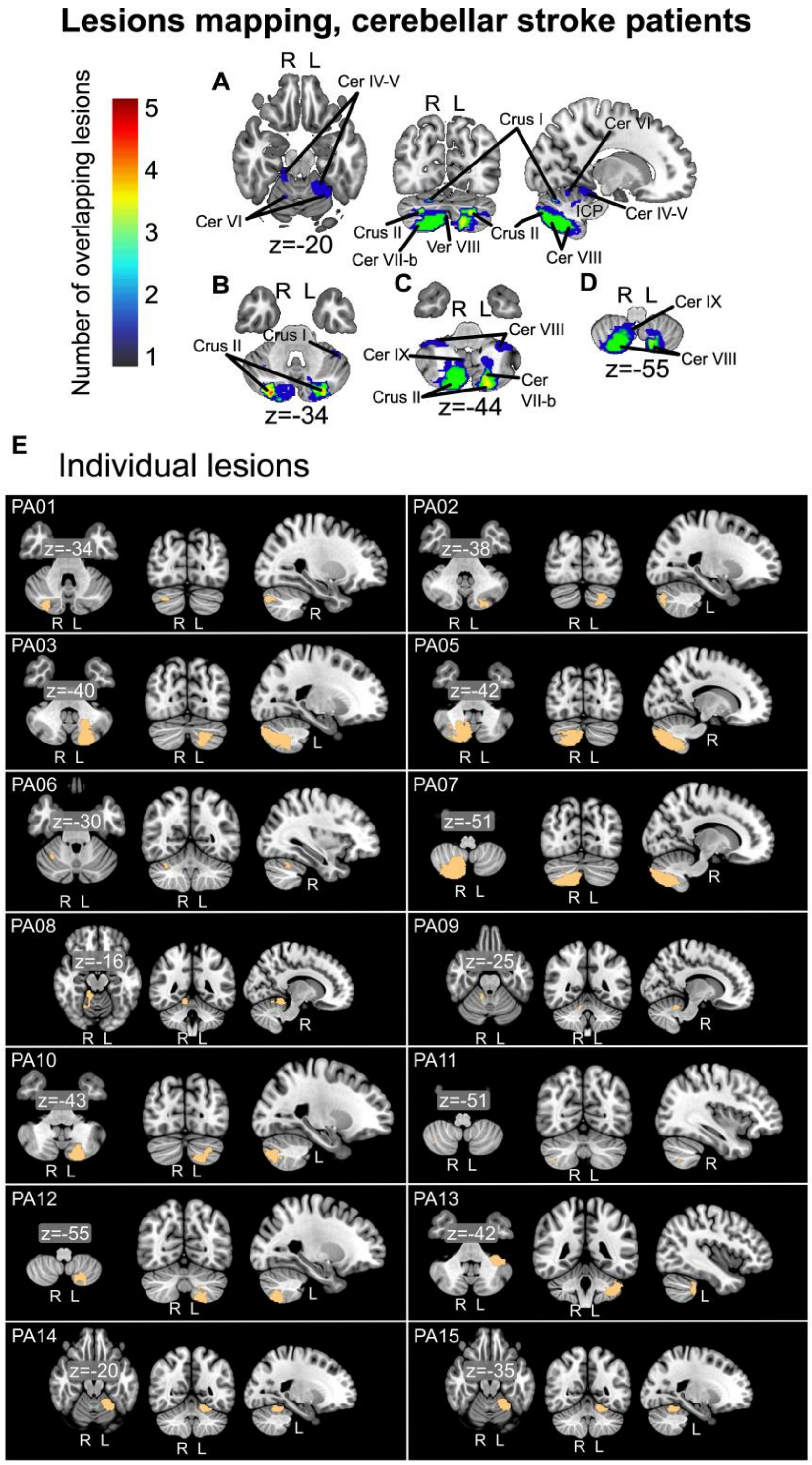

**Supplementary Figure 1: Mapping and overlaps of the lesions in the cerebellar stroke patients group. A**) Lesions are displayed in the MNI space on an axial, coronal and sagittal slice as well as at various Z-axis depth (**B**: MNI z=-34; **C**: MNI z=-44; **D**: MNI z=-55), and color reflects the number of patients having overlapping lesion territories, as indicated by the colorbar. **E**) Individual lesions, except for PA05 whose lesion was very hard to see, and which is located in the territory of the left superior cerebral artery. There are therefore 8 patients with left-lateralized lesions, and 7 with right-lateralized lesions. ICP: inferior cerebellar peduncle; Cer: cerebellar lobule; Ver: vermis. L/R: left/right hemisphere.

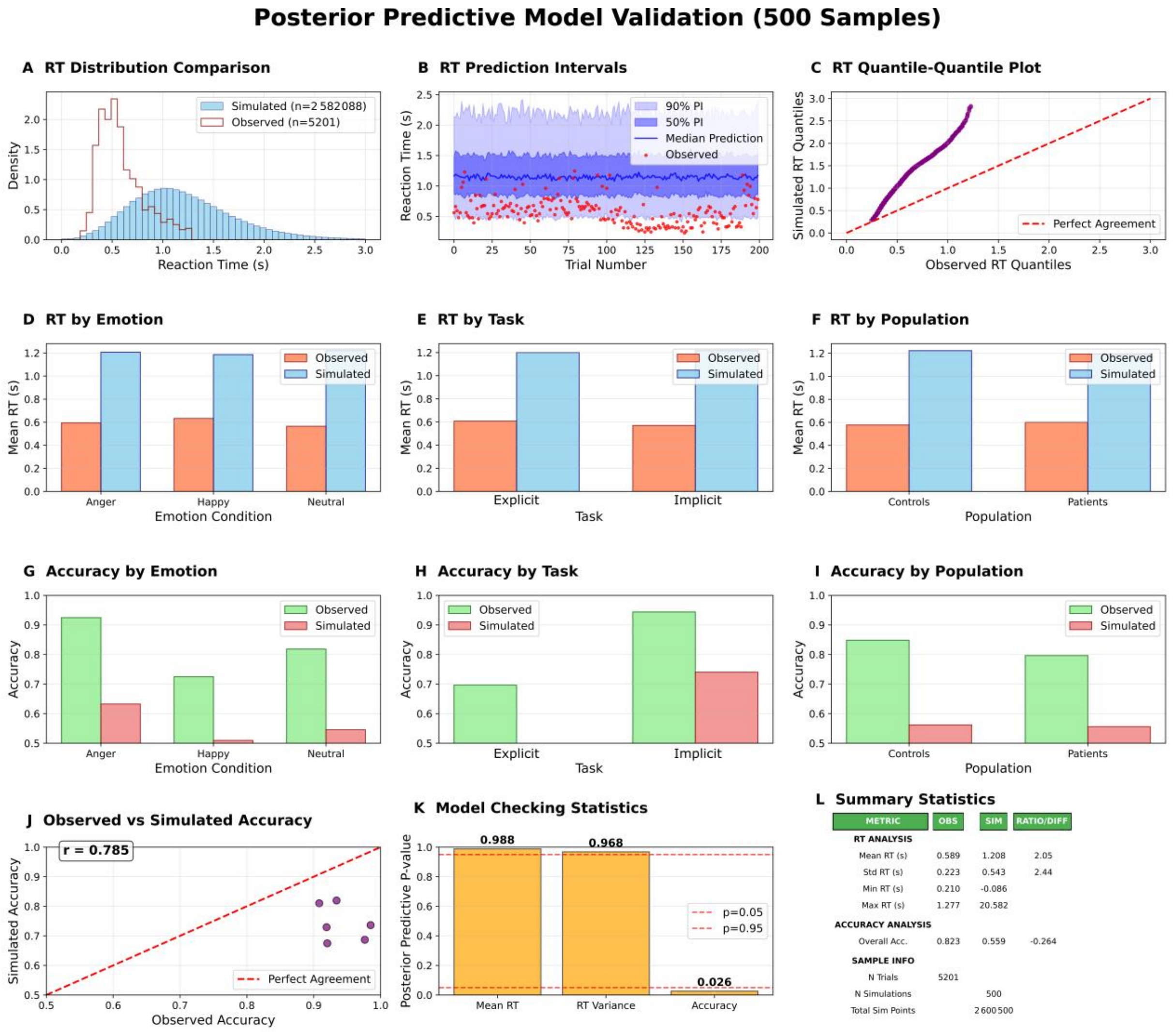

**Supplementary Figure 2: Posterior predictive model validation across 500 posterior samples.** (**A-C**) Reaction times distribution patterns. (**D-F**) Reaction times effects by factors. (**G-I**) Accuracy patterns by factors. (**J-L**) Overall model fit. RT: reaction times; diff: difference; N trials: number of trials; N simulations: number of simulations; Sim: simulated; Obs: observed; PI: prediction intervals.

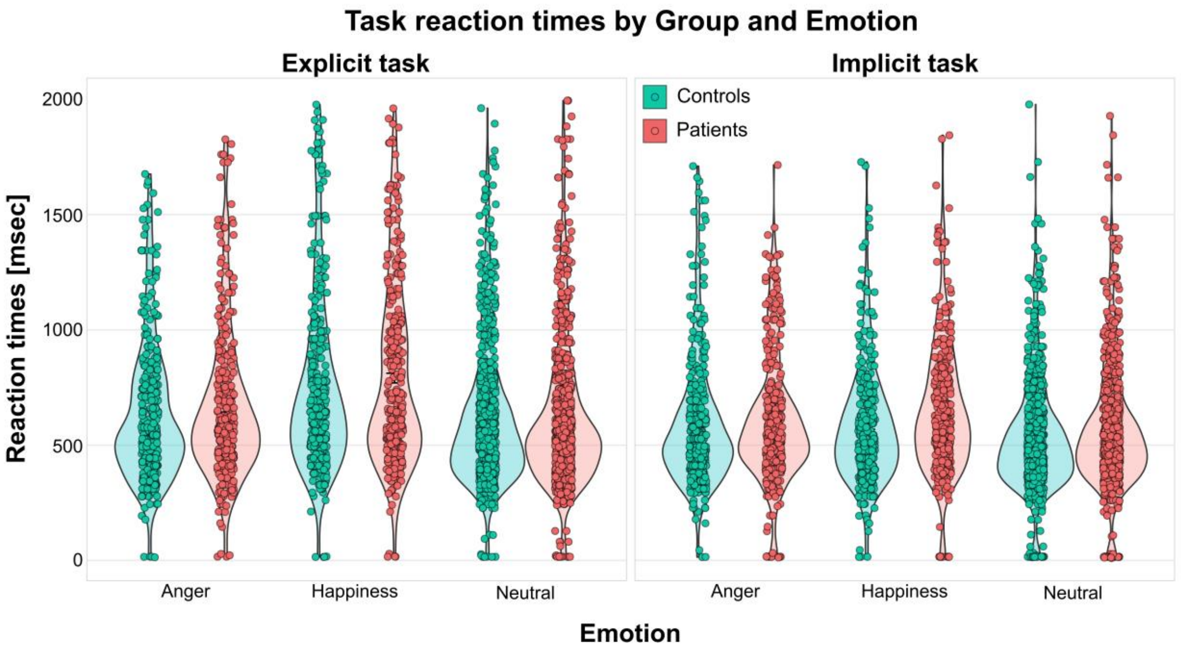

**Supplementary Figure 3. Voice categorization reaction times as a function of Emotion and Task, for both groups.** Response reaction times for voice categorization (*y* axis) for the explicit and implicit tasks in both control participants (green) and cerebellar stroke patients (coral) and according to each emotion (Anger/Happiness/Neutral, *x* axis). Bars represent the mean values, error bars the standard deviation from the mean, violins the shape of the distribution of the data and the points illustrate individual values, per trial and task, of each participant for each emotion.

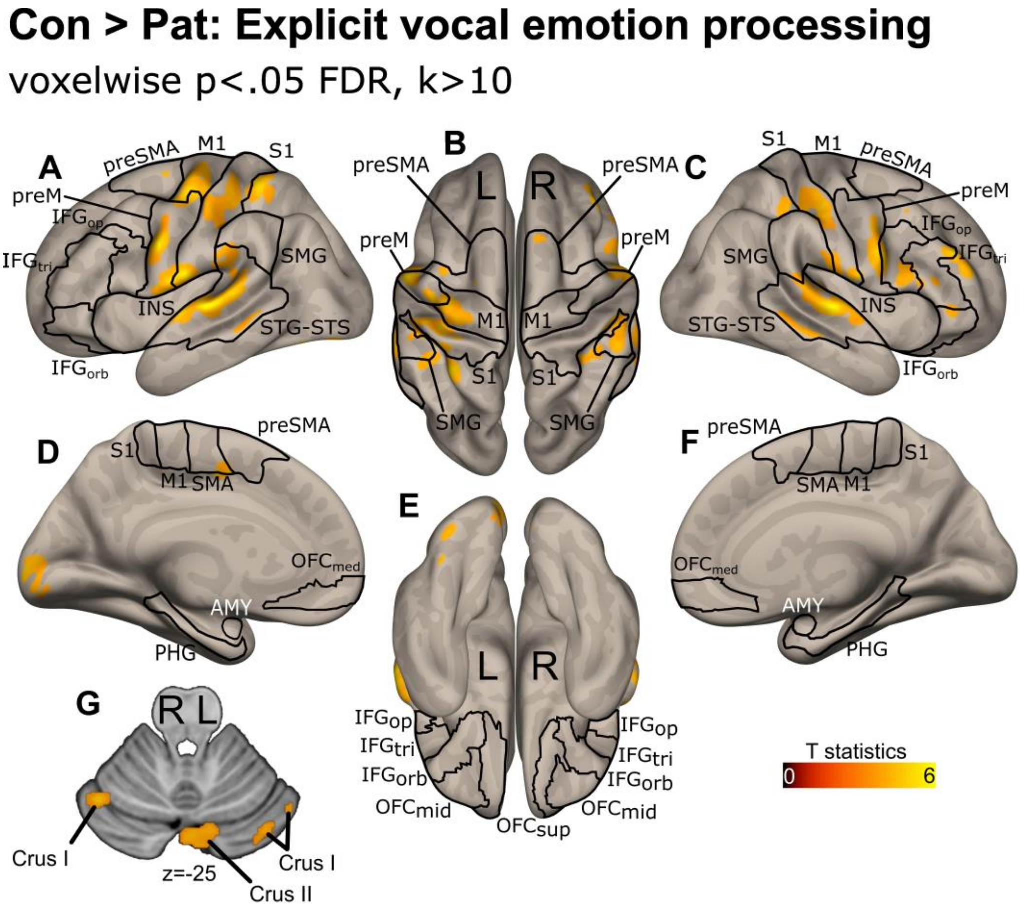

**Supplementary Figure 4. Neural activations when categorizing vocal emotion for the Explicit task, for controls compared to patients.** Sagittal (**A,C**), superior (**B**), medial (**D,F**), inferior (**E**) views in addition to an axial slice of the cerebellum (**G**) for the explicit categorization of emotional voices for controls (Con) compared to patients (Pat). Data corrected for multiple comparisons using voxelwise p<.05 FDR, with minimum cluster size of 10 or more voxels (k>10). The colorbar represents the voxelwise T value. IFG: inferior frontal gyrus; S1: primary somatosensory cortex; M1: primary motor cortex; preM: premotor cortex; SMG: supramarginal gyrus; STG/STS: superior temporal gyrus/sulcus; INS: insula; OFC: orbitofrontal cortex; AMY: amygdala; SMA: supplementary motor area; PHG: parahippocampal gyrus. Suffixes: op: *pars opercularis*; tri: *pars triangularis*; orb: *pars orbitalis*; med: medial; sup: superior. L/R: left/right hemisphere.

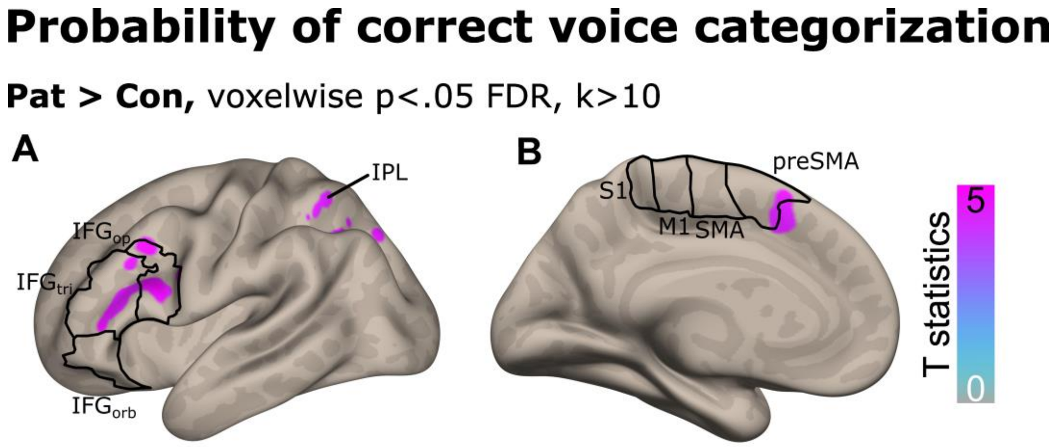

**Supplementary Figure 5: Neural correlates of the probability of correctly categorizing vocal emotion across tasks, by Population.** Sagittal view of the left hemisphere revealing lateral (**A**) and medial (**B**) brain correlates associated with the general probability of correct explicit and implicit categorization of emotional voices for patients (Pat) compared to control (Con) participants. Data corrected for multiple comparisons using voxelwise p<.05 FDR, with minimum cluster size of 10 or more voxels (k>10). The colorbar represents the voxelwise T value. IFG: inferior frontal gyrus; IPL: inferior parietal lobule; S1: primary somatosensory cortex; M1: primary motor cortex; SMA: supplementary motor area. Suffixes: op: *pars opercularis*; tri: *pars triangularis*; orb: *pars orbitalis*.

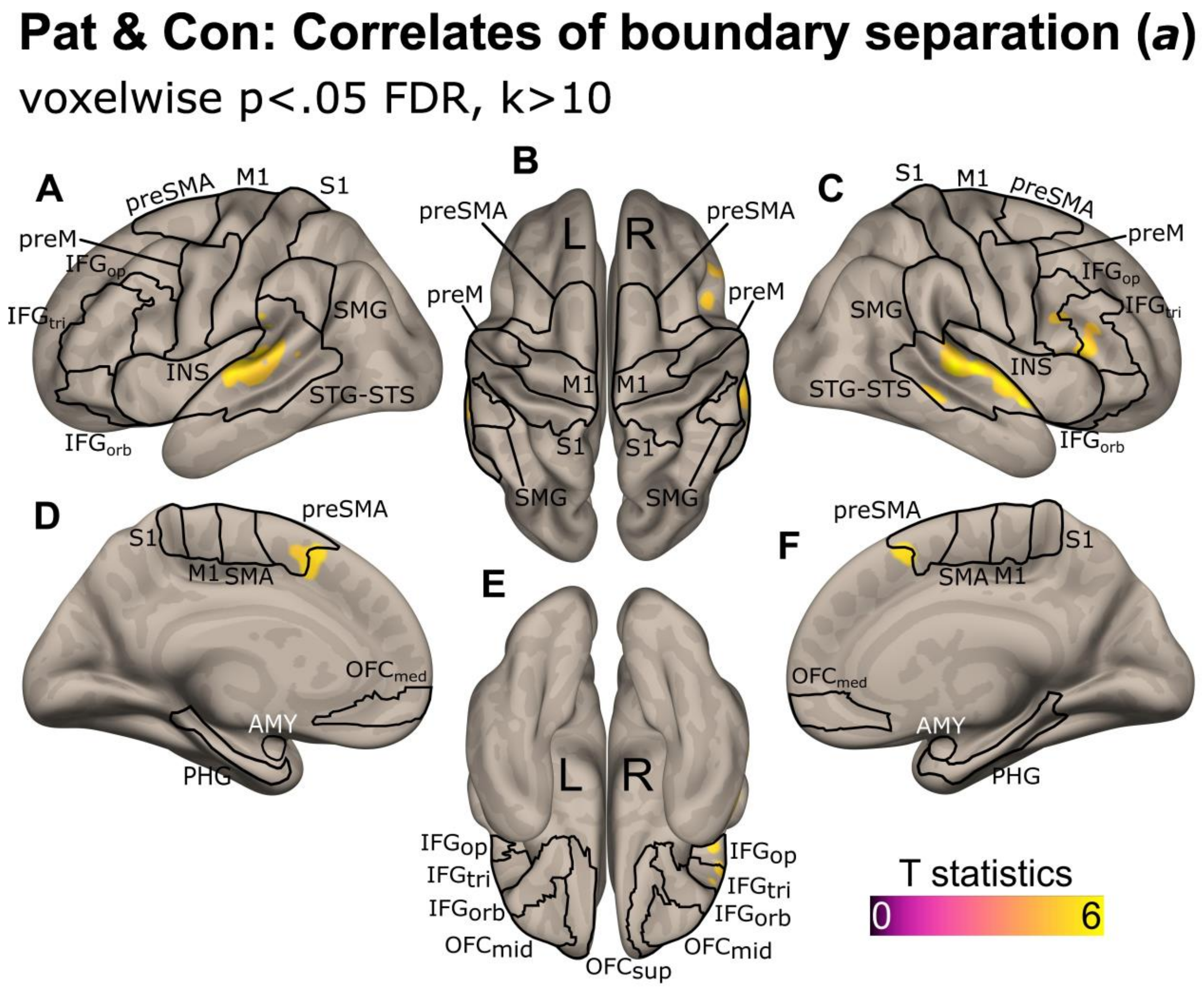

**Supplementary Figure 6: Neural correlates of boundary separation (*a*) following angle DDM modeling of the data, across Population.** Sagittal (**A,C**), superior (**B**), medial (**D,F**), inferior (**E**) views for decision caution, as represented by angle DDM boundary separation for Patients (Pat) and Controls (Con). Data corrected for multiple comparisons using voxelwise p<.05 FDR, with minimum cluster size of 10 or more voxels (k>10). The colorbar represents the voxelwise T value. IFG: inferior frontal gyrus; S1: primary somatosensory cortex; M1: primary motor cortex; preM: premotor cortex; SMG: supramarginal gyrus; STG/STS: superior temporal gyrus/sulcus; INS: insula; OFC: orbitofrontal cortex; AMY: amygdala; SMA: supplementary motor area; PHG: parahippocampal gyrus. Suffixes: op: *pars opercularis*; tri: *pars triangularis*; orb: *pars orbitalis*; med: medial; sup: superior. L/R: left/right hemisphere.

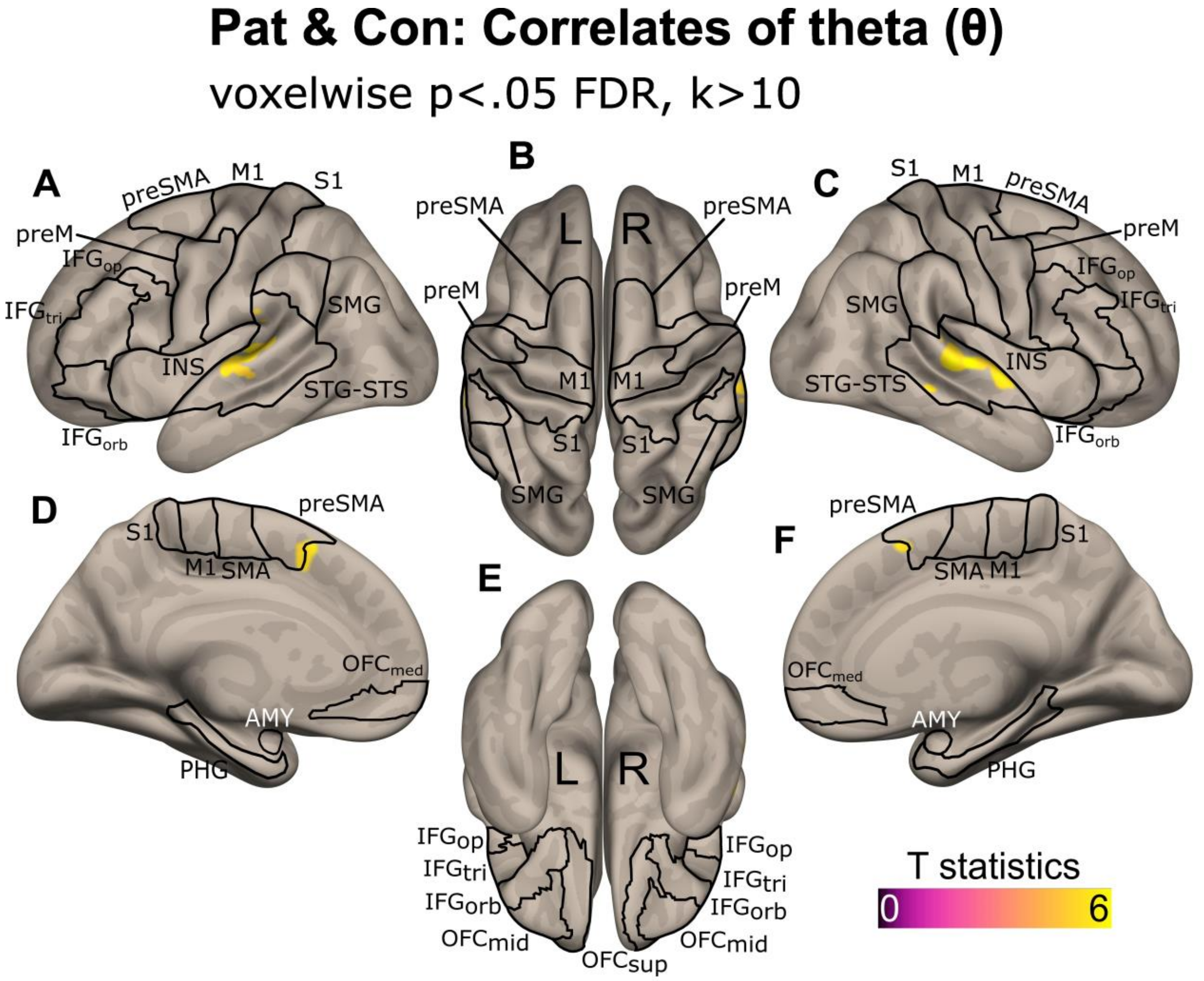

**Supplementary Figure 7: Neural correlates of theta (θ) following angle DDM modeling of the data, across Population.** Sagittal (**A,C**), superior (**B**), medial (**D,F**), inferior (**E**) views for attentional focus, as represented by angle DDM theta for Patients (Pat) and Controls (Con). Data corrected for multiple comparisons using voxelwise $p<.05$ FDR, with minimum cluster size of 10 or more voxels ($k>10$). The colorbar represents the voxelwise T value. IFG: inferior frontal gyrus; S1: primary somatosensory cortex; M1: primary motor cortex; preM: premotor cortex; SMG: supramarginal gyrus; STG/STS: superior temporal gyrus/sulcus; INS: insula; OFC: orbitofrontal cortex; AMY: amygdala; SMA: supplementary motor area; PHG: parahippocampal gyrus. Suffixes: op: *pars opercularis*; tri: *pars triangularis*; orb: *pars orbitalis*; med: medial; sup: superior. L/R: left/right hemisphere.

**Supplementary Table 1: Mean values for each dependent variable, for Task type, Emotion and Population factors.**

| Dependent variable : Accurate response probability, binarized ('0'=incorrect, '1'=correct), Artificially balanced model | | | | |
|---|---|---|---|---|
| *Task* | *Emotion* | *Population* | *BinaryResponse(mean)* | *BinaryResponse(SD)* |
| Explicit | Anger | Controls | 0.95 | 0.22 |
| Explicit | Anger | Patients | 0.89 | 0.31 |
| Explicit | Happy | Controls | 0.50 | 0.50 |
| Explicit | Happy | Patients | 0.45 | 0.49 |
| Explicit | Neutral | Controls | 0.69 | 0.46 |
| Explicit | Neutral | Patients | 0.65 | 0.48 |
| Implicit | Anger | Controls | 0.92 | 0.27 |
| Implicit | Anger | Patients | 0.91 | 0.28 |
| Implicit | Happy | Controls | 0.97 | 0.16 |
| Implicit | Happy | Patients | 0.92 | 0.27 |
| Implicit | Neutral | Controls | 0.99 | 0.11 |
| Implicit | Neutral | Patients | 0.92 | 0.28 |
| **Dependent variable : Reaction times [msec], Artificially balanced model** | | | | |
| *Task* | *Emotion* | *Population* | *ReactionTimes(mean)* | *ReactionTimes(SD)* |
| Explicit | Anger | Controls | 633 | 300 |
| Explicit | Anger | Patients | 660 | 323 |
| Explicit | Happy | Controls | 753 | 388 |
| Explicit | Happy | Patients | 791 | 391 |
| Explicit | Neutral | Controls | 625 | 312 |
| Explicit | Neutral | Patients | 624 | 332 |
| Implicit | Anger | Controls | 594 | 284 |
| Implicit | Anger | Patients | 605 | 275 |
| Implicit | Happy | Controls | 582 | 254 |
| Implicit | Happy | Patients | 646 | 286 |
| Implicit | Neutral | Controls | 527 | 246 |

| | | | | |
|---|---|---|---|---|
| Implicit | Neutral | Patients | 563 | 270 |
| **Dependent variable : Accurate response probability, binarized ('0'=incorrect, '1'=correct), Imbalanced model (most accurate design architecture)** | | | | |
| ***Task*** | ***Emotion*** | ***Population*** | ***BinaryResponse(mean)*** | ***BinaryResponse(SD)*** |
| Ex.Ang | Anger | Controls | 0.95 | 0.23 |
| Ex.Ang | Anger | Patients | 0.89 | 0.31 |
| Ex.Hap | Happy | Controls | 0.50 | 0.50 |
| Ex.Hap | Happy | Patients | 0.45 | 0.49 |
| **Ex.Ang** | **Neutral** | **Controls** | **0.89** | **0.31** |
| **Ex.Ang** | **Neutral** | **Patients** | **0.85** | **0.35** |
| **Ex.Hap** | **Neutral** | **Controls** | **0.49** | **0.50** |
| **Ex.Hap** | **Neutral** | **Patients** | **0.44** | **0.49** |
| Implicit | Anger | Controls | 0.92 | 0.27 |
| Implicit | Anger | Patients | 0.91 | 0.28 |
| Implicit | Happy | Controls | 0.97 | 0.16 |
| Implicit | Happy | Patients | 0.92 | 0.27 |
| Implicit | Neutral | Controls | 0.99 | 0.11 |
| Implicit | Neutral | Patients | 0.92 | 0.28 |

Ex.Ang: Explicit, anger-neutral block type; Ex.Hap : Explicit, happiness-neutral block type.
The splitting of the neutral voice stimuli according to the type of block is emphasized in bold.

**Supplementary Table 2: Comparisons for our Factors with drift diffusion 'angle' modeling of the data using Bayesian statistics.**

| | Factor | Comparison | Diff (avg) | CI-95 | P_Dir | Evidence |
|---|---|---|---|---|---|---|
| ***v*** | Pop | Patients vs Controls | -0.026 | [-0.50, +0.41] | 0.465 | Inconclusive |
| ***v*** | Task | Implicit vs Explicit | +1.191 | [+1.04, +1.34] | 1.000 | Very Strong |
| ***v*** | Emo | Happy vs Anger | -0.527 | [-0.95, -0.01] | 0.020 | Strong |
| ***v*** | Emo | Neutral vs Anger | -0.341 | [-0.60, +0.02] | 0.045 | Strong |
| ***v*** | Emo | Neutral vs Happy | +0.186 | [-0.44, +0.71] | 0.740 | Weak |
| ***a*** | Pop | Patients vs Controls | +0.014 | [-0.31, +0.40] | 0.505 | Inconclusive |
| ***a*** | Task | Implicit vs Explicit | +0.234 | [+0.16, +0.29] | 1.000 | Very Strong |
| ***a*** | Emo | Happy vs Anger | +0.052 | [-0.18, +0.36] | 0.665 | Weak |
| ***a*** | Emo | Neutral vs Anger | -0.106 | [-0.28, +0.22] | 0.175 | Weak |
| ***a*** | Emo | Neutral vs Happy | -0.158 | [-0.51, +0.23] | 0.175 | Weak |
| ***z*** | Pop | Patients vs Controls | +0.017 | [-0.28, +0.31] | 0.535 | Inconclusive |
| ***z*** | Task | Implicit vs Explicit | -0.009 | [-0.04, +0.02] | 0.300 | Inconclusive |
| ***z*** | Emo | Happy vs Anger | +0.017 | [-0.16, +0.15] | 0.660 | Weak |
| ***z*** | Emo | Neutral vs Anger | +0.017 | [-0.15, +0.20] | 0.630 | Weak |
| ***z*** | Emo | Neutral vs Happy | -0.001 | [-0.21, +0.24] | 0.480 | Weak |
| ***t*** | Pop | Patients vs Controls | +0.015 | [-0.42, +0.45] | 0.525 | Inconclusive |
| ***t*** | Task | Implicit vs Explicit | -0.036 | [-0.05, -0.02] | 0.000 | Very Strong |
| ***t*** | Emo | Happy vs Anger | +0.018 | [-0.19, +0.23] | 0.620 | Weak |
| ***t*** | Emo | Neutral vs Anger | +0.059 | [-0.12, +0.32] | 0.780 | Weak |
| ***t*** | Emo | Neutral vs Happy | +0.042 | [-0.22, +0.36] | 0.660 | Weak |
| **θ** | Pop | Patients vs Controls | +0.024 | [-0.31, +0.34] | 0.600 | Inconclusive |
| **θ** | Task | Implicit vs Explicit | +0.117 | [+0.05, +0.17] | 1.000 | Very Strong |
| **θ** | Emo | Happy vs Anger | +0.090 | [-0.12, +0.33] | 0.820 | Weak |
| **θ** | Emo | Neutral vs Anger | -0.071 | [-0.31, +0.14] | 0.220 | Weak |
| **θ** | Emo | Neutral vs Happy | -0.161 | [-0.45, +0.10] | 0.115 | Weak |

The Evidence column is based on the posterior probability values (P_Dir; including both positive and negative values): P_Dir > 0.95 or P_Dir < 0.05, "Strong"; P_Dir > 0.90 or P_Dir < 0.10, "Moderate"; P_Dir > 0.80 or P_Dir < 0.20, "Weak"; All other P_Dir values, "Inconclusive". *v*: drift rate; *a*: boundary separation; *z*: starting point; *t*: non-decision time; theta (θ): angle of collapsing boundaries; Diff: mean difference; CI-95: 95% confidence interval; P_Dir: value and direction of the posterior probability; Pop: Population factor; Emo: Emotion factor; Task: Task factor. Conclusive comparisons are colored in red.

**Supplementary Table 3: Interactions between our Factors with drift diffusion 'angle' modeling of the data using Bayesian statistics.**

| Two-way interactions | | | | | | | | |
|---|---|---|---|---|---|---|---|---|
| **P** | **F1** | **F2** | **Comparison** | **Diff (avg)** | **CI low** | **CI up** | **P_Dir** | **Evid** |
| v | Pop | Task | (Con×Expl-Con×Impl) - (Pat×Expl-Pat×Impl) | -9,25E-19 | -2,22E-16 | 2,22E-16 | 0,118 | W/Non |
| v | Pop | Emo | (Con×Hap-Con×Ang) - (Pat×Hap-Pat×Ang) | -2,87E-18 | -2,22E-16 | 2,22E-16 | 0,232 | W/Non |
| v | Pop | Emo | (Con×Neu-Con×Ang) - (Pat×Neu-Pat×Ang) | -3,29E-19 | -2,22E-16 | 2,22E-16 | 0,287 | W/Non |
| v | Task | Emo | (Impl×Hap-Impl×Ang) - (Expl×Hap-Expl×Ang) | -1,63E-17 | -2,50E-16 | 2,22E-16 | 0,230 | W/Non |
| v | Task | Emo | (Impl×Neu-Impl×Ang) - (Expl×Neu-Expl×Ang) | -2,00E-17 | -2,77E-16 | 2,22E-16 | 0,270 | W/Non |
| a | Pop | Task | (Con×Expl-Con×Impl) - (Pat×Expl-Pat×Impl) | 5,36E-18 | -4,44E-16 | 4,44E-16 | 0,258 | W/Non |
| a | Pop | Emo | (Con×Hap-Con×Ang) - (Pat×Hap-Pat×Ang) | -1,38E-17 | -4,44E-16 | 4,44E-16 | 0,217 | W/Non |
| a | Pop | Emo | (Con×Neu-Con×Ang) - (Pat×Neu-Pat×Ang) | -6,38E-18 | -4,44E-16 | 4,44E-16 | 0,227 | W/Non |
| a | Task | Emo | (Impl×Hap-Impl×Ang) - (Expl×Hap-Expl×Ang) | 1,44E-17 | -4,44E-16 | 4,44E-16 | 0,272 | W/Non |
| a | Task | Emo | (Impl×Neu-Impl×Ang) - (Expl×Neu-Expl×Ang) | -4,16E-18 | -4,44E-16 | 4,44E-16 | 0,235 | W/Non |
| z | Pop | Task | (Con×Expl-Con×Impl) - (Pat×Expl-Pat×Impl) | 3,25E-20 | -1,11E-16 | 1,11E-16 | 0,221 | W/Non |
| z | Pop | Emo | (Con×Hap-Con×Ang) - (Pat×Hap-Pat×Ang) | 2,62E-18 | -1,11E-16 | 1,11E-16 | 0,285 | W/Non |
| z | Pop | Emo | (Con×Neu-Con×Ang) - (Pat×Neu-Pat×Ang) | 3,41E-18 | -1,11E-16 | 1,11E-16 | 0,277 | W/Non |
| z | Task | Emo | (Impl×Hap-Impl×Ang) - (Expl×Hap-Expl×Ang) | -3,1E-18 | -1,11E-16 | 1,11E-16 | 0,200 | W/Non |
| z | Task | Emo | (Impl×Neu-Impl×Ang) - (Expl×Neu-Expl×Ang) | 6,07E-19 | -1,11E-16 | 1,11E-16 | 0,205 | W/Non |
| t | Pop | Task | (Con×Expl-Con×Impl) - (Pat×Expl-Pat×Impl) | 4,96E-18 | -1,11E-16 | 1,67E-16 | 0,248 | W/Non |
| t | Pop | Emo | (Con×Hap-Con×Ang) - (Pat×Hap-Pat×Ang) | -4,57E-18 | -1,11E-16 | 1,11E-16 | 0,240 | W/Non |

| | | | | | | | | |
|---|---|---|---|---|---|---|---|---|
| t | Pop | Emo | (Con×Neu-Con×Ang) - (Pat×Neu-Pat×Ang) | -7,1E-19 | -1,11E-16 | 1,11E-16 | 0,250 | W/Non |
| t | Task | Emo | (Impl×Hap-Impl×Ang) - (Expl×Hap-Expl×Ang) | 4,3E-18 | -1,11E-16 | 1,11E-16 | 0,235 | W/Non |
| t | Task | Emo | (Impl×Neu-Impl×Ang) - (Expl×Neu-Expl×Ang) | 2,54E-18 | -1,11E-16 | 1,11E-16 | 0,207 | W/Non |
| θ | Pop | Task | (Con×Expl-Con×Impl) - (Pat×Expl-Pat×Impl) | 4,04E-20 | -1,11E-16 | 1,11E-16 | 0,198 | W/Non |
| θ | Pop | Emo | (Con×Hap-Con×Ang) - (Pat×Hap-Pat×Ang) | 1,35E-18 | -1,11E-16 | 1,11E-16 | 0,170 | W/Non |
| θ | Pop | Emo | (Con×Neu-Con×Ang) - (Pat×Neu-Pat×Ang) | -7,63E-19 | -1,11E-16 | 1,11E-16 | 0,207 | W/Non |
| θ | Task | Emo | (Impl×Hap-Impl×Ang) - (Expl×Hap-Expl×Ang) | 4,05E-18 | -1,11E-16 | 1,11E-16 | 0,187 | W/Non |
| θ | Task | Emo | (Impl×Neu-Impl×Ang) - (Expl×Neu-Expl×Ang) | 1,24E-18 | -1,11E-16 | 8,39E-17 | 0,230 | W/Non |

**Three-way interactions**

| **P** | **F1** | **F2** | **F3** | **Comparison** | **Diff (avg)** | **CI low** | **CI up** | **P_Dir** | **Evid** |
|---|---|---|---|---|---|---|---|---|---|
| v | Pop | Task | Emo | [Con vs Pat] × [Expl vs Impl] × [Ang vs Hap] | -1,11E-16 | -0,14 | 0,13 | 0,48 | W/Non |
| v | Pop | Task | Emo | [Con vs Pat] × [Expl vs Impl] × [Ang vs Neu] | -4,44E-16 | -0,13 | 0,12 | 0,48 | W/Non |
| v | Pop | Task | Emo | [Con vs Pat] × [Expl vs Impl] × [Hap vs Neu] | -3,33E-16 | -0,13 | 0,13 | 0,50 | W/Non |
| a | Pop | Task | Emo | [Con vs Pat] × [Expl vs Impl] × [Ang vs Hap] | 4,44E-16 | -0,10 | 0,09 | 0,50 | W/Non |
| a | Pop | Task | Emo | [Con vs Pat] × [Expl vs Impl] × [Ang vs Neu] | -2,22E-16 | -0,10 | 0,09 | 0,49 | W/Non |
| a | Pop | Task | Emo | [Con vs Pat] × [Expl vs Impl] × [Hap vs Neu] | -6,66E-16 | -0,10 | 0,10 | 0,48 | W/Non |
| z | Pop | Task | Emo | [Con vs Pat] × [Expl vs Impl] × [Ang vs Hap] | 5,55E-17 | -0,06 | 0,06 | 0,51 | W/Non |
| z | Pop | Task | Emo | [Con vs Pat] × [Expl vs Impl] × [Ang vs Neu] | -5,55E-17 | -0,06 | 0,06 | 0,52 | W/Non |
| z | Pop | Task | Emo | [Con vs Pat] × [Expl vs Impl] × [Hap vs Neu] | -1,11E-16 | -0,06 | 0,07 | 0,51 | W/Non |
| t | Pop | Task | Emo | [Con vs Pat] × [Expl vs Impl] × [Ang vs Hap] | 0,00E+00 | -0,11 | 0,11 | 0,50 | W/Non |

| | | | | | | | | | |
|---|---|---|---|---|---|---|---|---|---|
| t | Pop | Task | Emo | [Con vs Pat] × [Expl vs Impl] × [Ang vs Neu] | -1,11E-16 | -0,11 | 0,10 | 0,48 | W/Non |
| t | Pop | Task | Emo | [Con vs Pat] × [Expl vs Impl] × [Hap vs Neu] | -1,11E-16 | -0,11 | 0,11 | 0,49 | W/Non |
| θ | Pop | Task | Emo | [Con vs Pat] × [Expl vs Impl] × [Ang vs Hap] | 0,00E+00 | -0,07 | 0,07 | 0,50 | W/Non |
| θ | Pop | Task | Emo | [Con vs Pat] × [Expl vs Impl] × [Ang vs Neu] | 0,00E+00 | -0,07 | 0,07 | 0,51 | W/Non |
| θ | Pop | Task | Emo | [Con vs Pat] × [Expl vs Impl] × [Hap vs Neu] | 0,00E+00 | -0,07 | 0,07 | 0,51 | W/Non |

The Evidence (Evid) column is based on the posterior probability values (P_Dir; including both positive and negative values): P_Dir > 0.95 or P_Dir < 0.05, "Strong"; P_Dir > 0.90 or P_Dir < 0.10, "Moderate"; P_Dir > 0.80 or P_Dir < 0.20, "Weak/None" (W/Non); All other P_Dir values, "Inconclusive". *v*: drift rate; *a*: boundary separation; *z*: starting point; *t*: non-decision time; theta (θ): angle of collapsing boundaries; Diff (avg): mean difference; CI low: lower bound of 95% confidence interval; CI up: upper bound of 95% confidence interval; P_Dir: value and direction of the posterior probability; Pop & F1: Population factor; Emo & F3: Emotion factor; Task & F2: Task factor; P: parameter estimates.

**Supplementary Table 4: Neural coordinates for the explicit task, comparing patients to controls (*p*<.05 FDR, k>10).**

| cluster size | T value | MNI X | MNI Y | MNI Z | Region | Hemisphere |
|---|---|---|---|---|---|---|
| 1077 | 7,31 | 4 | 60 | -10 | vmPFC | R |
| | 6,72 | 6 | 54 | -2 | OFC med | R |
| | 4,70 | 6 | 46 | -18 | OFC med | R |
| 238 | 6,41 | -4 | -30 | -18 | Int Caps | L |
| 427 | 5,96 | 26 | 10 | -18 | OLF | R |
| | 4,61 | 34 | 30 | -10 | IFGorb | R |
| | 4,06 | 20 | -4 | -16 | AMY | R |
| 295 | 5,85 | 2 | 32 | 26 | ACC | R |
| | 3,72 | -2 | 20 | 44 | preSMA | L |
| | 3,65 | 0 | 22 | 34 | CC mid | L |
| 265 | 5,34 | 30 | 28 | 42 | DLPFC | L |
| 428 | 5,19 | 44 | -62 | -12 | ITG post | L |
| | 4,79 | 52 | -52 | -14 | ITG post | L |
| | 4,26 | 44 | -46 | -12 | ITG post | L |
| 154 | 5,07 | -46 | -20 | -20 | ITG mid | L |
| 171 | 5,04 | 10 | -46 | 36 | PCC | R |
| | 3,79 | 4 | -36 | 50 | S1 | R |
| | 3,78 | 10 | -44 | 54 | Precuneus | R |
| 116 | 5,01 | -6 | -52 | -28 | Cer IX | L |
| 229 | 4,80 | 44 | -68 | 38 | AG | R |
| | 4,50 | 36 | -72 | 48 | AG | R |
| 100 | 4,76 | 52 | 8 | -32 | TP | R |
| | 3,85 | 52 | 12 | -24 | TP | R |
| 186 | 4,67 | 8 | -66 | 28 | Precuneus | R |
| | 3,51 | 0 | -64 | 38 | Precuneus | R |
| | 3,34 | -4 | -64 | 30 | Precuneus | L |
| 223 | 4,64 | 10 | -24 | -38 | Int Caps | R |
| 97 | 4,62 | 48 | -4 | -8 | STG mid | R |
| | 3,78 | 50 | -10 | -14 | STS mid | R |
| 67 | 4,61 | -8 | -42 | 32 | PCC | L |
| 434 | 4,50 | -30 | 18 | -8 | INS ant | L |
| | 4,17 | -48 | 0 | -12 | STG ant | L |
| | 4,12 | -22 | 2 | -14 | AMY | L |
| 53 | 4,46 | 32 | -36 | -28 | Cer IV-V | R |
| 124 | 4,44 | -2 | 24 | -8 | ACC subgenual | L |
| 104 | 4,44 | -34 | -74 | 38 | OC mid | L |
| 151 | 4,43 | -42 | -80 | -6 | OC inf | L |
| 143 | 4,42 | 14 | -48 | -54 | Cer IX | R |
| | 4,07 | 4 | -56 | -48 | Cer IX | R |
| | 3,71 | -6 | -52 | -56 | Cer IX | L |
| 58 | 4,36 | 48 | -6 | -34 | ITG ant | R |
| | 3,66 | 54 | -14 | -30 | ITG mid | R |

| | | | | | | |
|---|---|---|---|---|---|---|
| 20 | 4,11 | -36 | -32 | -24 | FFC | L |
| 190 | 4,10 | -26 | 32 | 38 | DLPFC | L |
| | 3,97 | -24 | 32 | 28 | DLPFC | L |
| | 3,61 | -24 | 46 | 30 | DLPFC | L |
| 73 | 4,10 | -50 | -58 | -16 | ITG post | L |
| 24 | 4,01 | 0 | -14 | 42 | CC mid | L |
| 44 | 3,98 | 18 | 30 | 52 | SFG | R |
| 28 | 3,95 | 46 | -24 | -18 | FFC | R |
| 107 | 3,91 | 26 | -86 | -8 | Lingual gyrus | R |
| 21 | 3,89 | -32 | -24 | -14 | Hipp | L |
| 43 | 3,85 | -20 | -58 | 2 | Lingual gyrus | L |
| 66 | 3,83 | 4 | 52 | 20 | ACC | R |
| 18 | 3,83 | -48 | 4 | -38 | ITG ant | L |
| 15 | 3,82 | 26 | -24 | -20 | PHG | R |
| 16 | 3,74 | -10 | 36 | 44 | SFG med | L |
| 15 | 3,73 | -18 | 20 | 6 | Caudate | L |
| 14 | 3,60 | -16 | -12 | -24 | PHG | L |
| 18 | 3,55 | -58 | -6 | -24 | ITG ant | L |
| 25 | 3,53 | -16 | 12 | -6 | Putamen | L |

vmPFC: ventromedial prefrontal cortex; OFC: orbitofrontal cortex; Int Caps: internal capsula; OLF: olfactory cortex; IFG: inferior frontal cortex; preSMA: pre-supplementary motor area; CC: cingulate cortex; ITG: inferior temporal cortex; PCC: posterior cingulate cortex; AG: angular gyrus; SFG: superior frontal gyrus; Hipp: hippocampus; PHG: parahippocampal gyrus; PCC: posterior cingulate cortex; TP: temporal pole; OC: occipital cortex; S1: primary somatosensory cortex; SFG: superior frontal gyrus; MTG: middle temporal gyrus; STG: superior temporal gyrus; FFA: fusiform face area; ACC: anterior cingulate cortex; DLPFC: dorso-lateral prefrontal cortex; AMY: amygdala; Cer: cerebellar lobule. Suffixes: ant, anterior; post, posterior; mid, mid; med, medial; orb, *pars orbitalis*.

**Supplementary Table 5: Neural coordinates for the implicit task, comparing patients to controls (*p*<.05 FDR, k>10).**

| cluster size | T value | MNI X | MNI Y | MNI Z | Region | Hemisphere |
|---|---|---|---|---|---|---|
| 648 | 6,55 | 34 | -70 | 44 | AG | R |
| 59 | 4,97 | 30 | 6 | 52 | DLPFC | R |
| 26 | 4,94 | 8 | -22 | -24 | Int Caps | R |
| 150 | 4,84 | 8 | -70 | 52 | Precuneus | R |
| | 4,33 | 6 | -64 | 60 | Precuneus | R |
| 136 | 4,66 | -22 | -76 | 46 | SPC | L |
| 104 | 4,55 | -42 | 4 | 38 | preM1 | L |
| | 4,40 | -48 | 4 | 46 | preM1 | L |
| 32 | 4,46 | 32 | 24 | 50 | MFG | R |
| 60 | 4,39 | -48 | -72 | -12 | OC inf | L |
| 63 | 4,38 | -4 | 20 | 46 | preSMA | L |
| | 3,70 | -4 | 16 | 36 | CC mid | L |
| 21 | 4,32 | 10 | -4 | 0 | Thalamus | R |
| 48 | 4,29 | -4 | -62 | -50 | Cer IX | L |
| | 3,87 | -8 | -52 | -52 | Cer IX | L |
| 42 | 4,24 | -54 | 2 | -12 | STG ant | L |
| 11 | 4,19 | -6 | -62 | -18 | Cer VI | L |
| 34 | 4,16 | -46 | 32 | 20 | IFGtri | L |
| 16 | 4,14 | -30 | 44 | 32 | DLPFC | L |
| 30 | 4,13 | -34 | 30 | 42 | DLPFC | L |
| 13 | 3,89 | -32 | 24 | 0 | INS ant | L |
| 40 | 3,82 | 8 | -72 | 32 | Precuneus | R |
| | 3,48 | 14 | -66 | 30 | Precuneus | R |
| 11 | 3,80 | -10 | -70 | 56 | Precuneus | L |
| 14 | 3,63 | 48 | 24 | -10 | IFGorb | R |

AG: angular gyrus; OC: occipital cortex; DLPFC: dorso-lateral prefrontal cortex; Int Caps: internal capsula; preSMA: pre supplementary motor area; SPC: superior parietal cortex; IFG: inferior frontal gyrus; MFG: middle frontal gyrus; STG: superior temporal gyrus; preM1: pre-primary motor cortex; INS: insula; Cer: cerebellar lobule; CC: cingulate cortex;OC: occipital cortex. Suffixes: ant, anterior; mid, mid; tri, *pars triangularis*; rob, *pars orbitalis*.

**Supplementary Table 6: Neural coordinates for the correlates of DDM drift rate (v) across tasks, for patients and controls (*p*<.05 FDR, k>10).**

| cluster size | T value | MNI X | MNI Y | MNI Z | Region | Hemisphere |
|---|---|---|---|---|---|---|
| 797 | 7,10 | 56 | 8 | -8 | TP | R |
| | 6,80 | 68 | -28 | 2 | STS mid | R |
| | 6,65 | 66 | -12 | 4 | STG mid | R |
| 952 | 6,68 | -54 | -20 | 2 | STG mid | L |
| | 6,25 | -64 | -22 | -2 | STSmid | L |
| | 5,93 | -64 | -14 | 0 | STG ant | L |
| 368 | 5,97 | 44 | 14 | 24 | IFGop | R |
| | 5,60 | 56 | 28 | 12 | IFGtri | R |
| | 5,06 | 48 | 28 | 16 | IFGtri | R |
| 192 | 5,66 | 2 | 18 | 54 | preSMA | R |
| | 4,67 | -4 | 12 | 54 | SMA | L |
| | 4,61 | 12 | 26 | 52 | preSMA | R |
| 69 | 5,66 | -46 | 20 | -2 | IFGtri | L |
| | 4,18 | -36 | 20 | -2 | INS ant | L |
| | 4,12 | -44 | 10 | 2 | IFO | L |
| 226 | 5,56 | -48 | 18 | 22 | IFGop | L |
| | 4,30 | -42 | 20 | 32 | IFGop | L |
| | 4,27 | -50 | 22 | 34 | DLPFC | L |
| 25 | 4,96 | -40 | 2 | -34 | ITG | L |
| 18 | 4,34 | 4 | 30 | 38 | CC mid | R |
| 13 | 4,29 | 48 | 8 | 40 | MFG | R |
| 16 | 3,95 | -40 | 40 | 0 | IFGtri | L |

TP: temporal pole; STS: superior temporal sulcus; IFO: inferior frontal operculum; preSMA: pre-supplementary motor area; SMA: supplementary motor area; IFG: inferior frontal gyrus; DLPFC: dorsolateral prefrontal cortex; CC: cingulate cortex; MFG: middle frontal gyrus; ITG: inferior temporal gyrus; STG: superior temporal gyrus; INS: insula. Suffixes: ant, anterior; mid, mid; tri, *pars triangularis*; op, *pars opercularis*.

**Supplementary Table 7: R-hat convergence values for the modeling of our data through DDM angle, per parameter**

| Parameter | N Parameters | R-hat Range | Mean R-hat |
|---|---|---|---|
| Boundary separation ($a$) | 22 | 1.0 - 1.0007 | 1,0002 |
| Drift Rate ($v$) | 22 | 0.9999 - 1.0006 | 1,0002 |
| Non-decision Time ($t$) | 22 | 1.0 - 1.001 | 1,0004 |
| Starting Point ($z$) | 22 | 1.0 - 1.0005 | 1,0003 |
| Theta (θ) | 22 | 1.0 - 1.001 | 1,0004 |

N Parameters: number of parameters, namely the combination of each parameter with the factors of the global formula described and explained in the methods. Common R-hat values and their meaning for convergence are the following: R-hat ≤ 1.01: excellent; R-hat ≤ 1.05: good; R-hat > 1.05: poor; R-hat > 1.10: very poor convergence.

**Supplementary Table 8: Descriptive statistics for drift diffusion 'angle' modeling of the data**

| | *v* estimate | *a* estimate | *z* estimate | *t* estimate | θ estimate |
|---|---|---|---|---|---|
| **Trial count** | 5201 | 5201 | 5201 | 5201 | 5201 |
| **Mean posterior value** | 1,69 | 1,10 | 0,45 | 0,15 | 0,55 |
| **Std** | 0,67 | 0,13 | 0,01 | 0,03 | 0,07 |
| **Min** | 0,60 | 0,88 | 0,43 | 0,11 | 0,45 |
| **25%** | 1,04 | 0,99 | 0,43 | 0,12 | 0,52 |
| **50%** | 1,79 | 1,11 | 0,45 | 0,15 | 0,57 |
| **75%** | 2,24 | 1,23 | 0,46 | 0,18 | 0,64 |
| **Max** | 2,64 | 1,28 | 0,47 | 0,22 | 0,67 |

Parameters (columns) are the posterior mean value. *v*: drift rate; *a*: boundary separation; *z*: starting point; *t*: non-decision time; theta (θ): angle of collapsing boundaries.
